# Cooperation and cheating orchestrate *Vibrio* assemblages and polymicrobial synergy in oysters infected with OsHV-1 virus

**DOI:** 10.1101/2023.02.11.528104

**Authors:** Daniel Oyanedel, Arnaud Lagorce, Maxime Bruto, Philippe Haffner, Amandine Morot, Yann Dorant, Sébastien de La Forest Divonne, François Delavat, Nicolas Inguimbert, Caroline Montagnani, Benjamin Morga, Eve Toulza, Cristian Chaparro, Jean-Michel Escoubas, Yannick Labreuche, Yannick Gueguen, Jeremie Vidal-Dupiol, Julien de Lorgeril, Bruno Petton, Lionel Degremont, Delphine Tourbiez, Léa-Lou Pimparé, Marc Leroy, Océane Romatif, Juliette Pouzadoux, Guillaume Mitta, Frédérique Le Roux, Guillaume M. Charrière, Marie-Agnès Travers, Delphine Destoumieux-Garzón

## Abstract

Polymicrobial diseases significantly impact the health of humans and animals but remain understudied in natural systems. We recently described the Pacific Oyster Mortality Syndrome (POMS), a polymicrobial disease that impacts oyster production and is prevalent worldwide. Analysis of POMS-infected oysters on the French North Atlantic coast revealed that the disease involves co-infection with the endemic ostreid herpesvirus 1 (OsHV-1) and virulent bacterial species such as *Vibrio crassostreae*. However, it is unknown whether consistent *Vibrio* populations are associated with POMS in different regions, how *Vibrio* contribute to POMS, and how they interact with the OsHV-1 virus during pathogenesis.

We resolved the *Vibrio* population structure in oysters from a Mediterranean ecosystem and investigated their functions in POMS development. We find that *Vibrio harveyi* and *Vibrio rotiferianus* are the predominant species found in OsHV-1-diseased oysters and show that OsHV-1 is necessary to reproduce the partition of the *Vibrio* community observed in the field. By characterizing the interspecific interactions between OsHV-1, *V. harveyi* and *V. rotiferianus*, we find that only *V. harveyi* synergizes with OsHV-1. When co-infected, OsHV-1 and *V. harveyi* behave cooperatively by promoting mutual growth and accelerating oyster death. *V. harveyi* showed high virulence potential in oysters and dampened host cellular defenses, making oysters a more favorable niche for microbe colonization. We next investigated the interactions underlying the co-occurrence of diverse *Vibrio* species in diseased oysters. We found that *V. harveyi* harbors genes responsible for the biosynthesis and uptake of a key siderophore called vibrioferrin. This important resource promotes the growth of *V. rotiferianus*, a cheater that efficiently colonizes oysters during POMS without costly investment in host manipulation nor metabolite sharing.

By connecting field-based approaches, laboratory infection assays and functional genomics, we have uncovered a web of interdependencies that shape the structure and function of the POMS pathobiota. We showed that cooperative behaviors contribute to synergy between bacterial and viral co-infecting partners. Additional cheating behaviors further shape the polymicrobial consortium. Controlling such behaviors or countering their effects opens new avenues for mitigating polymicrobial diseases.

## Introduction

A number of polymicrobial diseases impact human and animal species [1]. They are defined as diseases that result from infections by multiple pathogens [2]. Complex microbe communities that form a cohesive entity with the potential to cause disease in polymicrobial diseases can be referred to as a “pathobiota” [3]. Within pathobiota, microbes synergize to cause disease: their interactions enhance disease progression compared to infection with the single microbes [1]. Fatal polymicrobial synergy has been reported between viruses and bacteria including influenza A virus and *Streptococcus* pneumonia [4, 5] or Human Immunodeficiency Virus (HIV) and *Mycobacterium tuberculosis* [6-8]. Synergy has also been reported between viruses such as *Herpes* virus simplex and HIV [9, 10]. Pathobiota show cell-level cooperative capacities that include the production of public goods (e.g. shared metabolites), division of labor, resource transport, and creation and maintenance of the extracellular environment, as in other examples of multicellular organization [11]. Dissecting the interactions within pathobiota is needed to understand and possibly control disease establishment, progression, and symptoms. Theoretical models have been developed in response to these challenges to predict what microbial behaviors are favored during such complex interactions [12, 13]. In addition, a series of animal models (both vertebrates and invertebrates) have been used to mimic polymicrobial diseases and validate theoretical assumptions [1]. Still, natural pathobiota remain poorly explored.

We recently described a typical example of a polymicrobial disease, the Pacific Oyster Mortality Syndrome (POMS). This disease is caused by the Ostreid herpesvirus OsHV-1 and opportunistic bacteria [14] and has devastating consequences for the aquaculture of *Crassostrea gigas* oysters worldwide. The bacterial genera colonizing oysters during POMS are conserved across environments suggesting functional complementarity within the pathobiota [15]. Members of the *Vibrionaceae* family are the best characterized bacteria in the POMS pathobiota [16-20]. Several *Vibrio* species have been shown to have virulence functions in this disease [19, 21]. *Vibrio crassostreae* (Splendidus clade) uses cytotoxicity and other mechanisms to evade oyster cellular immune responses, leading to systemic infection [19, 21]. This virulent *Vibrio* species is highly prevalent in OsHV-1-infected oysters on the French Atlantic coast, often associated with other *Vibrio* species from the superclade Splendidus [17]. A number of studies suggest that *Vibrio* species found in OsHV-1-infected oysters vary worldwide [22-24]. These species include *Vibrio harveyi* (superclade Harveyi). A strain of *Vibrio harveyi* was isolated in 2003 from *C. gigas* oyster spat during a mortality episode in the Thau lagoon (Mediterranean sea) and is pathogenic to oysters [25]. However, we still know little about *Vibrio harveyi* in POMS and whether *V. crassostreae* or other populations colonize OsHV-1-infected oysters in different ecosystems. In addition, how *Vibrio* colonizes oysters and how *Vibrio* interacts with the OsHV-1 virus during pathogenesis remains unclear.

Here we performed an integrative study of POMS in a Mediterranean ecosystem, combining field analysis of the *Vibrio* population structure in OsHV-1-infected oysters with validation of this polymicrobial assembly in mesocosm experiments. We find that two species of the Harveyi clade – namely *Vibrio harveyi* and *Vibrio rotiferianus* – are prevalent in diseased oysters in a major Mediterranean area used for oyster farming, the Thau lagoon. Using mesocosm experiments, we characterized the complex interactions between *Vibrio* species and the OsHV-1 virus as well as between the different *Vibrio* species that assemble in OsHV-1-infected oysters. Our data indicate that OsHV-1 infection favors stable colonization by *V. harveyi* and *V. rotiferianus* but not other *Vibrio* of the Harveyi clade. Polymicrobial synergy, including mutual growth promotion and accelerated disease progression, was measured between OsHV-1 and *V. harveyi*. We next tested the contribution of each partner to the interaction. A series of functional assays, including gene knockouts, indicate that strains of *V. harveyi* are cytotoxic to immune cells and produce siderophores. Our results uncover multiple interdependencies within the POMS pathobiota leading to polymicrobial synergy and accelerated disease progression. We find that initial infection with OsHV-1 shapes *Vibrio* assemblages within the host and favors colonization by *V. harveyi*. This colonization promotes the growth of OsHV-1 and *V. rotiferianus* in oysters by dampening host defenses and by producing vibrioferrin, a key siderophore required for *Vibrio* growth in iron-poor environments.

## Materials and methods

### Oyster and seawater sampling

Juvenile *C. gigas* (pathogen-free diploid oysters produced in hatchery, 6 months old) were immersed in the Thau lagoon (Occitanie, France) and sampled at four timepoints between October 2015 and March 2017. Sampling coincided with mortality events and in the absence of any observed mortality (see Table S1 and S2 for details). 40L of seawater was collected and size-fractionated by sequential filtration (from > 60 μm to 0.2 μm) as described by Bruto et al. [17].

### Bacterial sampling

Oysters and large seawater particle fractions (> 60 μm) were ground up with Ultra-Turrax (IKA) and 100 μL was plated on *Vibrio* selective media (thiosulfate-citrate-bile salts sucrose agar, TCBS). Filters of 5, 1 and 0.22 μm porosity were directly placed on TCBS agar and incubated at 20 °C for 2 days. About 100 colonies per sample were randomly picked then re-streaked first on TCBS and then on Zobell agar (4 g.L^-1^ bactopeptone, 1 g.L^-1^ yeast extract and 15 g.L^-1^ agar in sterile seawater, pH 7.4). Stock cultures were stored at −80 °C in Zobell containing 15% glycerol (v/v). For subsequent molecular analyses, all isolates were grown overnight at 20 °C in liquid Zobell medium and bacterial DNA was extracted using the Nucleospin tissue kit following the manufacturer’s instructions (Macherey-Nagel).

### Population structure analysis

Isolates from October 2015 were genotyped by partial *hsp60* sequencing [26, 27] (Table S3) generating a total of 437 *hsp60* sequences [28]. *hsp60* sequence ambiguities were corrected using 4 peaks and Seaview software (http://nucleobytes.com/index.php/4peaks; [29]) and received a taxonomic affiliation if the best BLAST-hit displayed an identity greater than 95% with a type-strain. Fisher-exact tests were performed with a 2×2 contingency table using the computing environment R [30] for statistical validation of the ecological preferences of populations and the distribution of bacterial populations in oyster tissues and seawater. Significance was assessed using p-value ⩽ 0.05.

### MLSA genotyping

To validate *hsp60* sequence-based taxonomic assignments to the Harveyi clade, 3 additional protein-coding genes were sequenced (*rctB, topA* and *mreB*). First, Harveyi isolates were screened by PCR using rctB-F/rctB-R primers designed to specifically hybridize to Harveyi-related *rctB* gene sequences (Table S3). The PCR program was: 2 min at 95 °C; 30 cycles of 30 sec at 95 °C, 1 min at 53 °C and 1.45 min at 72 °C; 5 min at 72 °C. As a result of this analysis, 143 *rctB+* isolates were considered to belong to the Harveyi clade. Then *topA* and *mreB* sequences were amplified using VtopA400F/VtopA1200R and VmreB12F/VmreB999R primers, respectively [31, 32], Table S3). All three genes were amplified using the Gotaq G2 flexi polymerase (Promega) following the manufacturer’s instructions and sequenced using the reverse PCR primer at GATC Biotech. Next, *hsp60, rctB, topA* and *mreB* sequences were aligned with 8 Harveyi superclade type strains using Muscle [33]. Alignments were concatenated with Seaview [29]. Phylogenetic trees for each marker were reconstructed with RAxML using a GTR model of evolution and Gamma law of rate heterogeneity. Bootstrap values were calculated for 100 replicates. All other options were left at default values.

### Bacterial growth conditions

Bacteria were grown for 18 h under shaking at 20°C in Zobell liquid medium (4 g.L^-1^ bactopeptone, 1 g.L^-1^ yeast extract in sterile seawater, pH 7.4) or LB broth adjusted to 0.5M NaCl unless otherwise stated. When necessary, antibiotics were added (Trimethoprim Trim 10 μg/mL or Chloramphenicol Cm 10 μg/mL).

### Virulence potential of *Vibrio* strains

To test virulence potential, *Vibrio* were grown under shaking at 20 °C for 18 h in Zobell liquid medium before adjustment to OD_600_ = 0.7. A volume of 40 μL was injected intramuscularly into 20 specific pathogen-free (SPF) juvenile *C. gigas* oysters [16] previously anesthetized in hexahydrate MgCl_2_ (50 g.L^−1^, 100 oysters/liter). An injection of *V. crassostreae* J2-9 (virulent strain), *V. tasmaniensis* LMG20012^T^ (non-virulent strain) or sterile filtered seawater (negative control) were used as controls. After injection, animals were transferred to aquaria (20 oysters per 1 L aquarium) containing 400 mL of aerated seawater at 20 °C and kept under static conditions. Mortalities were recorded 24 h post injection.

### *In vitro* cytotoxicity assays

Hemocytes were plated in 96 well-plates (2 × 10^5^ cells/well) as previously published [34]. After 1 h, plasma was removed and 5 μg/μL Sytox Green (Molecular Probes) diluted in 200 μL sterile seawater was added to each well. Washed *Vibrios* that had been opsonized in plasma for 1 h were then added to the wells at an MOI of 50:1. Sytox Green fluorescence was monitored (ƛex 480 nm/ƛem 550 nm) for 15 h using a TECAN microplate reader. Maximum cytolysis was determined by adding 0.1% Triton X-100 to hemocytes. Statistical analysis was performed using one-way ANOVA and a post-hoc Tukey test for pairwise comparison of maximum cytolysis values for each condition. Significance was assessed using p-value ⩽ 0.05.

### Fluorescence microscopy

Hemocytes were plated onto glass coverslips in a 24-well plate to obtain monolayers of 5 × 10^5^ cells per well. Adherent hemocytes were exposed to GFP or mCherry-expressing Washed *Vibrios* that had been opsonized in plasma for 1 h, were then added to the wells at a MOI of 50:1. *Vibrios* (Table S4) were added at a multiplicity of infection of 50:1, as in [34]. Binding of bacteria to hemocytes was synchronized by centrifugation for 5 min at 400 *g*. After a 2 h incubation, the cell monolayers (coverslips from bottom of the wells) were fixed with 4 % paraformaldehyde for 15 min. Coverslips were then washed in PBS and stained with 0.25 μg.mL^-1^ DAPI (Sigma) and 0.5 μg.ml^-1^ Phalloidin-TRITC or FITC (Sigma). Fluorescence imaging was performed using a Zeiss Axioimager fluorescence microscope and a Zeiss 63× Plan-Apo 1.4 oil objective equipped with a Zeiss MRC black and white camera for image acquisition.

### Experimental infection in mesocosm

For experimental infections, a biparental family of oyster (*C. gigas*) spat was produced at the Ifremer facilities in La Tremblade (Charente-Maritime, France). This family was selected for its susceptibility to OsHV-1 infection. Spawn occurred in June 2017, and larval and spat cultures were performed as described by Dégremont et al. [35] and Azéma et al. [36]. All growth steps involved filtered and UV-treated seawater. Prior to the experiment, spat were acclimated via a constant flow of filtered and UV-treated seawater enriched in phytoplankton (*Skeletonema costatum, Isochrysis galbana*, and *Tetraselmis suecica*) in 120 L tanks at 19°C for at least 2 weeks. Oysters (10 months, 4 cm) were infected with OsHV-1 virus, *Vibrio*, or both. Microorganisms were prepared as follows.

*Viral inoculation -* For **Design 1** (Fig. S1), seawater containing OsHV-1 virions was produced to infect pathogen-free juvenile oysters. Briefly, 90 donor oysters were anesthetized, and their adductor muscles were injected with 100 μl of 0.2 μm filtrated viral suspension (10^8^ genomic units mL^-1^). These donor oysters were then placed in a 40 L tank for 24 h. Virion release into the seawater was quantified by qPCR. This OsHV-1-contaminated seawater was used to fill tanks for the different experimental conditions. At day 0, 10 recipient oysters were placed in tanks containing 2 L of OsHV-1-contaminated seawater and were sampled at 4, 24, and 48 h; 15 additional recipient oysters were placed in tanks with 3 L of contaminated seawater to track mortalities daily. An identical design was used for control tanks, in which clean seawater was used instead of OsHV-1-contaminated water. For **Design 2** (Fig. S1), 250 donor oysters were injected with a filtrated viral suspension as described above. After 24 h, 250 recipient oysters were placed in contact with donor oysters in a 40 L-tank. After another 18 h (day 0), recipient oysters were transferred into clean seawater for mortality recording (10 animals in 0.5 L) or for sampling (30 animals in 1.5 L).

*Bacterial inoculation –* Seawater tanks were inoculated with bacteria on day 0 (final concentration of 10^7^ CFU/mL). Briefly, bacterial cultures (Zobell broth, 20°C, 18h) were centrifuged at 1500 x g for 10 min. Bacterial pellets were rinsed and resuspended in sterile seawater and the concentration was adjusted to OD_600_ = 1 (10^9^ CFU/mL). The bacterial concentration was confirmed by conventional dilution plating and CFU counting on Zobell agar.

At each sampling time point, oysters were sampled together with 100 mL of seawater. The oyster flesh was removed from the shell, snap-frozen in liquid nitrogen and stored at -80°C. 30 mL of seawater were filtered (0.2 μm pore size) and filters were stored at -80 °C. For tissue grinding, individual frozen oysters were shaken for 30 s inside a stainless steel cylinder containing a stainless-steel ball cooled in liquid nitrogen in a Retsch MM400 mixer mill. The pulverized tissue was transferred to a 2 mL screw-capped tube and stored at -80°C until further processing.

### Nucleic acid extraction

Total DNA was extracted from either 20 mg of frozen oyster tissue-powder, 25 mg frozen oyster tissue, a pellet from 1 mL of stationary phase bacterial cultures, or a 0.2 μm filter using the Nucleospin tissue DNA extraction kit (Macherey Nagel, ref: 740952.250) with a modified protocol. Briefly, samples were added to a 2 mL screw-capped tube containing Zirconium beads, lysis buffer, and proteinase K and shaken for 12 min at a frequency of 35 cycles/s in a Retsch MM400 mixer mill at room temperature and then incubated for 1h 30 min at 56 °C. The samples were then treated with RNase for 5 min at 20 °C and then 10 min at 70 °C. The following purification steps were carried out according to manufacturer’s recommendations. Total RNA was extracted from 20 mg of frozen oyster tissue-powder using Direct-zol RNA extraction kit (Zymo research). In an extra step, the aqueous phase was recovered from the TRIzol reagent prior to column purification as described by [37]; the following steps were carried out as recommended by the manufacturer. Nucleic acid concentration and purity was assayed using a Nanodrop ND-10000 spectrophotometer (Thermo Scientific) and RNA integrity analyzed by capillary electrophoresis on the BioAnalyzer 2100 system (Agilent).

### *Vibrio* genome sequencing and assembly

Individual genomic libraries were prepared from 1 ng of bacterial DNA at the Bio-Environment platform (University of Perpignan) using the Nextera XT DNA Library Prep Kit (Illumina) according to the manufacturer’s instructions. The quality of the libraries was checked using High Sensitivity DNA chip (Agilent) on a Bioanalyzer. Pooled libraries were sequenced in 2×150 paired-end mode on a NextSeq 550 instrument (Illumina). Reads were assembled *de novo* using Spades software. Computational prediction of coding sequences together with functional assignments and comparative genomics were performed using the MaGe MicroScope [38]. The genome sequence assemblies have been deposited in the European Nucleotide Archive (ENA) at EMBL-EBI under project accession no. PRJEB49488 (Table S6).

### OsHV-1 detection and quantification

The ground oyster flesh (1.5 mL) was centrifuged for 10 min at 4°C and 2,200 g and genomic DNA was extracted from 50 μL of supernatant using phenol:chloroform:isoamyl alcohol (25:24:1) and isopropanol precipitation. Detection and quantification of OsHV-1 μVar DNA was performed using quantitative PCR targeting a predicted DNA polymerase catalytic subunit (DP) using OsHVDPFor/OsHVDPRev primers (Table S3) [39] using the protocol previously described by [37]. All amplification reactions were performed in duplicate using a Roche LightCycler 480 Real-Time thermocycler (qPHD-Montpellier GenomiX platform, Montpellier University). Oyster flesh samples exhibiting a viral load greater than 100 genome units per ng of total DNA (GU/ng) were considered to be infected by OsHV-1.

### *Vibrio* quantification

16S rDNA sequences were used to quantify total *Vibrio* present in oysters by extracting 25 ng of DNA from tissue or crude extracts from seawater (see above). Amplification reactions were carried out in duplicate, in a total volume of 20 μl on Mx3005 Thermocyclers (Agilent) using Brilliant III Ultra-Fast SyberGreen Master Mix (Agilent), and 567F and 680R primers at 0.3 μM, [40], Table S3. Absolute quantification of *Vibrio* genomes in oyster samples was estimated using standards from 10^2^ to 10^9^ genome copies of *Vibrio* (see supplementary material and methods).

*V. harveyi-V. rotiferianus* and *V. owensii-V. jasicida* were quantified based on 25 ng of DNA extracted from tissue or crude extracts from seawater (see above) through detection of a specific chemotaxis protein and *ompA*, respectively (Table S3). As described above, amplification reactions were performed in duplicate using a Roche LightCycler 480 Real-Time thermocycler, SYBR Green I Master mix (Roche) and primers at 0.3 and 0.2 μM f.c. respectively. For absolute quantification, standard curves of known concentration of *V. harveyi, V. rotiferianus, V. jasicida, V. owensii* genomes were used (supplementary material and methods).

### *rctB* metabarcoding

Locus-specific PCR primers, including Illumina overhang adaptors, were designed to amplify a 573 bp region of the *rctB* gene in all our *Harveyi* strains, (rctB-Fw-I and rctB-Rv-I primers, Table S3). PCR analysis of total DNA extracted from oysters (N=60) used the high fidelity Q5 polymerase (New England Biolabs) in a total volume of 50 μL under the following conditions: 98 °C for 25 s followed by 35 cycles of 98 °C for 10 sec, 51 °C for 25 sec and 72 °C for 30 sec. Final extension was performed at 72 °C for 2 minutes. Presence of the 573 bp amplicon was validated by 1.5 % gel electrophoresis. Libraries were constructed with the Two-Step amplicon sequencing approach using Illumina dual indexes (ref. 15044223) and sequenced on a MiSeq instrument to produce paired end reads 2×300 bp, by the GenSeq platform, University of Montpellier (ISEM), France. Sequencing data were processed using the SAMBA pipeline v3.0.1. [41]. All bioinformatics processes used the next-generation microbiome bioinformatics platform QIIME 2 [42] (version 2020.2) and grouped sequences in ASV (Amplicon Sequence Variants) using DADA2 v1.14 [43]. The resulting ASVs were annotated against an in-house database containing the Harveyi *rctB* sequences and filtered for low abundance ASVs to limit the prevalence of putative artifacts due to sequencing errors. To do this, we only retained ASVs showing at least four reads in at least four samples. Statistical analyses were performed with R [30] using the R packages Phyloseq v1.38.0 [44] and Vegan v2.6-2 [45]. Principal coordinate analyses (PCoA) based on Bray-Curtis distances at each kinetic point were used to assess variation in the composition of Harveyi communities. Putative differences between groups were assessed by statistical analyses (Permutational Multivariate Analysis of Variance - PERMANOVA) using the function adonis2 implemented in vegan [45]. Finally, we used DESeq2 v1.36.0 and STAMP software [46] to identify ASVs with significant variation in abundance.

### Mutagenesis

Deletion of *pvuA1* (THOG05_v1_100041) and *pvuA2* (THOG05_v1_100042) in *V. rotiferianus* Th15_O_G05 was achieved through double homologous recombination between the pLP12 suicide plasmid (Table S8) and the bacterial chromosome [47]. Briefly, two fragments of around 800 bp flanking the target region were amplified, assembled by GeneArt, and cloned into the pLP12 plasmid [48]. The suicide plasmid (named pAM010) was transferred by conjugation between an *Escherichia coli* β3914 donor [49] and *V. rotiferianus* Th15_O_G05 recipient using a triparental mating procedure (Table S4-5). The first and second recombination events leading to pAM010 integration and elimination were selected following a recently published method [47]. Mutants were screened by PCR using primers del-pvuA1-A2-OG05-F and del-pvuA1-A2-OG05-R (Table S3). A *V. rotiferianus* Th15_O_G05 mutant strain deleted for the *pvuA1*-*2* genes was stored in glycerol at -80 °C (strain *V. rotiferianus* Th15_O_G05 Δ*pvuA1-2*).

### *Vibrio* growth in iron-depleted media and rescue

Growth experiments in iron-depleted medium were performed with *V. rotiferianus* Th15_O_G05 and its Δ*pvuA1-2* isogenic derivative. Isolates were grown overnight at room temperature in Artificial Sterile Seawater (ASW: NaCl 40 mM, KCl 20 mM, MgSO_4_ 5 mM, CaCl_2_ 2 mM) with 0.3 % (wt/vol) casamino acids and vitamins (0.1 μg/L vitamin B12, 2 μg/L biotin, 5 μg/L calcium pantothenate, 2 μg/L folic acid, 5 μg/L nicotinamide, 10 μg/L pyridoxin hydrochloride, 5 μg/L riboflavin, 5 mg/L thiamin hydrochloride). Cultures were pelleted (2 min at 15,000 g) and washed in ASW. Cells were then inoculated (1:100) into minimal media with (iron-poor) or without (iron-replete) the iron-specific chelator 2,2’-bipyridin (150 μM). Bacteria were grown in 96-well plates with orbital shaking in a Tecan microplate reader (Infinite M200) at 25 °C for 24 h. The OD_600_ was recorded at 30 min intervals. Rescue was performed either by adding freeze-drying concentrated *V. harveyi* Th15_O_G11 cell-free supernatant or by adding synthetic vibrioferrin (8.7 to 70 μM) to cultures. Vibrioferrin was synthesized according to Takeuchi et al. [50](Fig. S9). NMR and HRMS spectroscopic data confirmed the structure (Fig. S10) and were consistent with the literature [50].

## Results

### Specific *Vibrio* populations assemble in oysters infected by OsHV-1 virus

We asked whether *Vibrio* populations that naturally assemble in oysters infected by OsHV-1 were conserved across farming environments. With this objective in mind, we performed a field experiment in the Thau lagoon (South of France). *Vibrio* populations have been suggested to be distinct in this Mediterranean ecosystem from those previously found in the Atlantic (bay of Brest, northwest of France) [25]. We performed our study during an episode of oyster mortality. Specific pathogen-free (SPF) juvenile oysters were immersed in September 2015 in the Thau lagoon, which hosts significant oyster farming activity (352 acres). Oysters tested positive for OsHV-1 after one month (October 2015), indicating an ongoing episode of POMS (Table S1). We characterized the population structure of the *Vibrio* isolated from a pool of oysters and from the surrounding seawater. A total of 472 isolates were sampled on *Vibrio*-selective medium from infected oyster tissues and from the water column (Table S2). Partial *hsp60* sequences were obtained for 437 isolates. Assignment to the *Vibrio* genus was confirmed for 304 sequences (67.8 %), which exhibited ≥ 95 % identity with the *hsp60* sequence from a *Vibrio* type-strain (Table S2)., Isolates with *hsp60* sequence identity below this threshold were not included in the study. We observed contrasting population structures in oyster tissues and in the water column. Oyster tissues were dominated by 3 populations of the *Vibrio* Harveyi super clade (*V. harveyi, V. rotiferianus* and *V. owensii*), representing 54/55 isolates (Fig. 1A). The water column showed a higher diversity of *Vibrio* with only 53/249 isolates (21%) falling into the Harveyi super clade (Fig. 1A, Fig. S2). These water column isolates were dominated by *V. jasicida* (39/53) (Fig. 1A). Taxonomic assignments of Harveyi-related isolates were confirmed by multi-locus sequence analysis (MLSA) phylogeny based on 4 *Vibrio* genes (*hsp60, rctB, topA* and *mreB*) (Fig. 1B) (Lagorce, 2022). A total of 101 isolates could be assigned to *V. harveyi* (n=63), *V. rotiferianus* (n=17), *V. jasicida* (n=15) and *V. owensii* (n=6) (Fig. 1B). Among them, *V. harveyi* and *V. rotiferianus* showed a positive association with oyster tissues (Fisher exact test, *p* < 0.001) whereas *V. jasicida* was almost exclusively associated with the water column (Fisher exact test, *p* < 0.001) (Fig. 1B). The high relative abundance of *V. harveyi* in POMS-diseased oysters from the Thau lagoon was confirmed by three subsequent samplings in 2016-2017: *V. harveyi* was isolated during but not outside of POMS episodes, almost exclusively from oysters infected with OsHV-1 (Table S1). This result contrasts with the preferential association of the species *V. crassostreae* with POMS-diseased oysters on the French Atlantic coast [17].

**Figure 1.**
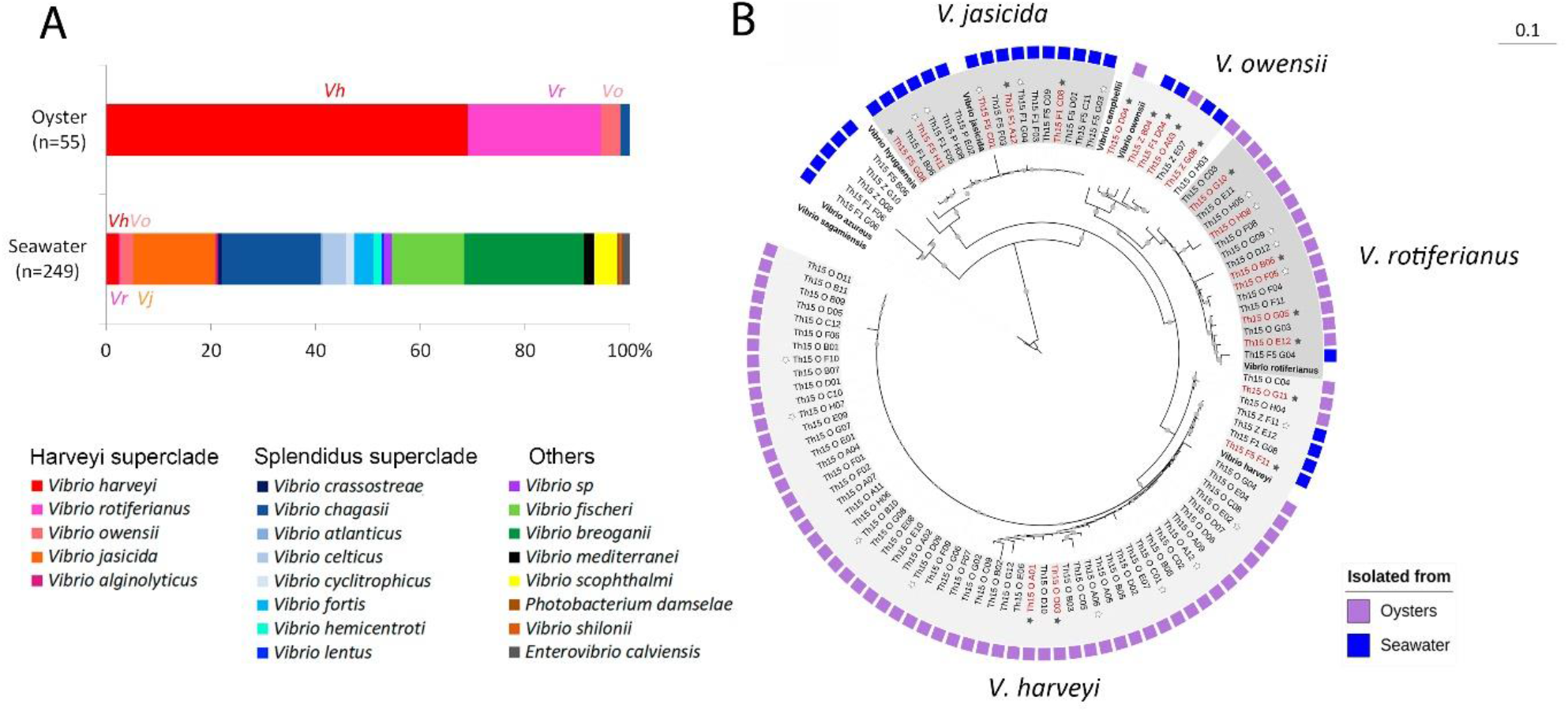
*Vibrio harveyi* and *Vibrio rotiferianus* are the most prevalent *Vibrio* species in OsHV-1-infected oysters. The population structure of *Vibrionaceae* was determined in seawater and oyster flesh during an episode of POMS (Thau lagoon, October 2015). **(A)** shows the distribution (%) of *Vibrio* isolated from oysters and seawater. A total of 304 isolates whose *hsp60* sequence displayed ≥ 95% identity with a *Vibrio* type strains were included. A high prevalence of the Harveyi super-clade (*V. harveyi* –*Vh, V. owensii –Vo, V. jasicida –Vj* and *V. rotiferianus* –*Vr*) is observed in oysters. **(B)** shows the high prevalence of *V. harveyi* and *V. rotiferianus* in oysters after taxonomic affiliation shown in A was validated by an MLST analysis. Four marker genes (*hsp60, rctB, topA* and *mreB)* were used. A phylogenetic tree was constructed with the concatenated sequences of the 4 markers. The following reference strains were used in the analysis: *V. azureus* NBRC 104587, *V. campbellii* CAIM 519, *V. harveyi* NBRC 15634, *V. hyugaensis* 090810a, *V. jasicida* CAIM 1864, *V. owensii* CAIM 1854, *V. rotiferianus* CAIM 577, *V. sagamiensis* NBRC 104589. *Vibrio sagamiensis* type strain was used as an outgroup to root the tree. Bootstrap values (>50%) are indicated by circles on branches. Color boxes indicate whether a strain was isolated from seawater (blue boxes) or from oysters (purple boxes). For details on seawater column fractionation, see Fig. S2. Stars indicate strains whose genome was sequenced in this study, and filled stars those used for comparative genomics. Red indicates strains used in mesocosm experiments (Fig. 2).

### Preferential association of *V. harveyi* and *V. rotiferianus* with OsHV-1-infected oysters

As a number of interdependent environmental variables (*e*.*g*. temperature, salinity, season, viral infection) may influence *Vibrio* assemblages in oysters in the field, we next tested how *Vibrio* populations partition and the role of OsHV-1 in this partitioning. To achieve this aim, we used mesocosms, which allow controlled infection experiments of oysters through natural routes with microorganisms relevant to POMS. We developed a synthetic *Vibrio* community composed of a mixture of 20 isolates from the Harveyi super clade in the presence (VO) or absence (V) of OsHV-1 (Fig. 2A). *Vibrio* representative of the 4 populations isolated from the Thau lagoon were used: *V. harveyi* and *V. rotiferianus* (positively associated with oysters), *V. jasicida* (negatively associated with oysters) and *V. owensii* (neutral). In parallel, oysters were exposed to OsHV-1 virus only (O) or were kept in tanks devoid of introduced pathogens (control). Oysters were collected in the first 48 h, before mortalities occurred. We first examined which *Vibrio* populations colonized oysters in the presence/absence of the OsHV-1 virus by comparing the V and VO conditions. To discriminate between the four populations introduced in the mesocosm, we developed an amplicon sequencing method based on the *rctB* polymorphic gene (Fig. S3). In the absence of OsHV-1, the Harveyi-related population assemblage remained stable in oysters over time, as shown by *rctB*-barcoding (Fig. S5). In contrast, co-infection with OsHV-1 had a significant effect on the structure of the assemblage, as observed after 48 h (Permutational multivariate analysis of variance, *p* = 0.001) (Fig. 2B, Fig. S4-S5). The species *V. harveyi* and *V. rotiferianus* were significantly enriched in oyster flesh in the presence of OsHV-1 (*p* < 0.05, Welch’s t-test, p-value corrected with Benjamini-Hochberg FDR) (Fig. 2C). In contrast, *V. owensi* was equally abundant in the presence/absence of OsHV-1, and *V. jascida* was more abundant in oyster flesh in the absence of OsHV-1 (*p* < 0.001) (Fig. 2C). The positive effect of OsHV-1 on oyster colonization by *V. harveyi/V. rotiferianus* but not *V. jasicida/V. owensii* at 48 h was confirmed by qPCR monitoring of pathogen loads (mutiple t test, *p* < 0.01) (Fig. S4). Altogether, our experimental results show (i) that OsHV-1 is necessary to reproduce the distribution of *Vibrio* community observed in the field during a POMS episode, and (ii) that only *V. harveyi* and *V. rotiferianus* efficiently colonize OsHV1-infected oysters.

**Figure 2.**
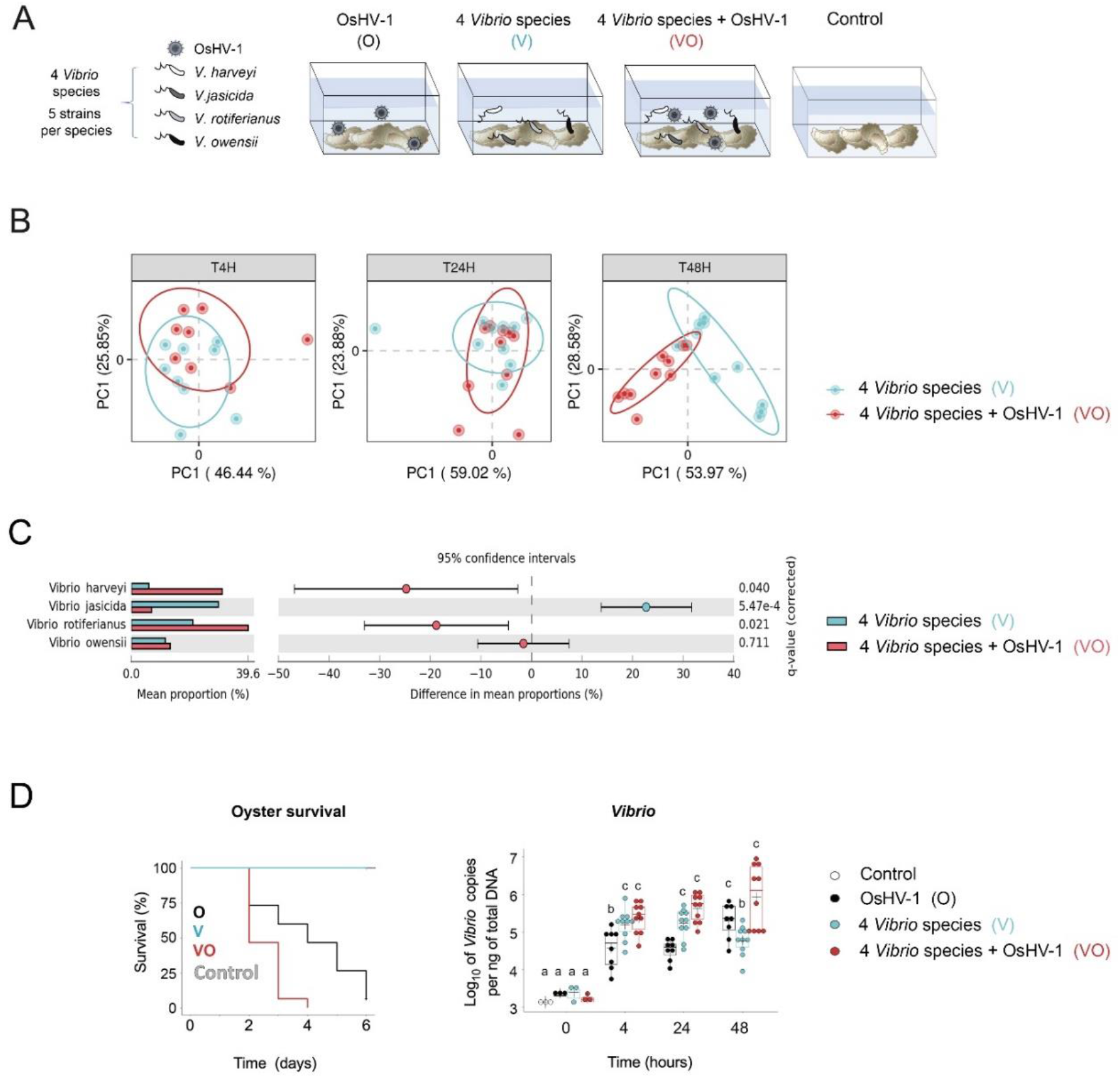
OsHV-1 synergizes with *Vibrio harveyi* and/or *Vibrio rotiferianus* to kill oysters. **(A) Mesocosm experiment to test OsHV-1 synergy with *Vibrio* species (Design 1)**. Specific pathogen-free oysters were placed in contact with seawater containing OsHV-1 (10^8^ genomic units mL^-1^) or *Vibrio* (10^7^ CFU.mL^-1^) or both during 6 days at 20°C. (O): seawater containing only OsHV-1. (V): 4 species of the Harveyi clade (V; *V. harveyi, V. rotiferianus, V. owensii, V. jasicida*), (VO): both OsHV-1 and *Vibrio*, (C): Controls not exposed to pathogen. **(B) Monitoring of the OsHV-1-induced dysbiosis by *rctB-*barcoding**. Principal Component Analysis (PCoA) ordination plots of Bray-Curtis dissimilarities for the Harveyi-related community associated with oysters. PCoA results are depicted for each time point (i.e. 4, 24 and 48 h). Each dot represents the *Vibrio* microbiome of one oyster. Colors refer to the experimental condition (*blue* for V, *red* for VO). Ellipses represent the 95% confidence intervals for each group. PERMANOVA between the two experimental conditions and the three time-points indicates significant differences p = 0.001 (see Fig S5). Statistical differences between V and VO conditions are observed at 48 h. **(C) Colonization of diseased oysters by *V. harveyi* and *V. rotiferianus* at 48 h**. The left barplot represents the mean proportion of the species *V. harveyi, V. rotiferianus, V. owensii, V. jasicida* in oysters from the V and VO conditions at T=48h. The right dot plot represents the difference in mean proportion of each species by STAMPS analysis. Statistical differences were obtained from Welch’s t-test (p-value corrected with Benjamini-Hochberg FDR). **(D) Synergy between OsHV-1 and *V. harveyi*/*V. rotiferianus***. Left panel shows a significant increase in the mortality rate for oysters exposed to both Harveyi and OsHV-1 (in red) compared to OsHV-1 only (in black) (Kaplan-Meier survival curves, log-rank test, p = 0.0018). Right panel shows that oyster colonization by *Vibrio* is favored by OsHV-1 (Kruskal-Wallis test, p value < 0.001).

### OsHV-1 virus synergizes with *V. harveyi* and/or *V. rotiferianus* in oyster mortality

We next tested the effect of the *Vibrio*/OsHV-1 interaction on POMS progression. We used the same synthetic community of *Vibrio* (Fig. 2A) to monitor oyster mortality and pathogen loads under different experimental conditions. Mortalities were only observed in tanks containing the OsHV-1 virus, indicating that the synthetic *Vibrio* community alone (V) was not lethal to oysters through natural infection routes. Mortalities started at day 2 in tanks containing OsHV-1 (both O and VO conditions). Still, mortalities progressed significantly more rapidly in oyster tanks containing both OsHV-1 and the synthetic *Vibrio* community (VO) with 90% mortalities in 3 days as opposed to 6 days for oysters exposed to OsHV-1 only (O) (Kaplan-Meier survival curves, log-rank test, *p* = 0.0018) (Fig. 2D, left panel). Therefore, introducing the synthetic *Vibrio* community accelerated the OsHV-1-induced disease. As expected from our previous study [14], the total *Vibrio* load increased constantly over 48 h in OsHV-1-infected oysters (O). Remarkably, the increase in *Vibrio* load was significantly higher in oysters exposed to both the *Vibrio* community and OsHV-1 (VO) than in oysters exposed to *Vibrio* only (O) at 48h, *i*.*e*. at the onset of mortality (Kruskal-Wallis test, *p* < 0.001) (Fig. 2D, right panel), with a significant contribution of *V. harveyi* and *V. rotiferianus* (Fig. S4). In oysters exposed to *Vibrio* only (V), *i*.*e*. oysters that did not die, *Vibrio* colonization tended to be transient with a peak between 4 h-24 h (Fig. 2D, right panel). Altogether, this indicates that in the absence of OsHV-1, oysters tolerate transiently high loads of the synthetic *Vibrio* community but *Vibrio* colonization is not stable and ultimately decreases without causing mortality. In contrast, OsHV-1 infection favors the proliferation and persistent colonization of *Vibrio* such as *V. harveyi* and *V. rotiferianus*, which exacerbate pathogenesis, an effect not seen with *V. jascida* and *V. owensii*. These data show that OsHV-1 and specific populations of *Vibrio* act in synergy to accelerate oyster death.

### *V. harveyi* and OsHV-1 reciprocally promote inside-host growth

To get insight into the synergistic process, we next tested whether OsHV-1 and strains of *V. harveyi* or *V. rotiferianus* affect one another’s growth in co-infections. To facilitate pathogen monitoring oysters were exposed to OsHV-1 and fluorescent *Vibrio* strains representing each population (Fig. 3A). Infection with OsHV-1 only was used as a control (Fig. 3A). As in the previous mesocosm experiment (Fig 2), live oysters were sampled at 0, 4, 24 and 48 h, before the onset of mortalities (Fig. S6) to monitor pathogen loads in every individual. First, we compared *V. harveyi* and *V. rotiferianus* colonization in oysters infected with OsHV-1. Only *V. harveyi* had the ability to colonize OsHV-1-infected oysters efficiently. Indeed, *V. harveyi* remained present at high doses (10^5^ to 5 × 10^6^ copies/ng of DNA) in live oyster tissues throughout the time course. In contrast, *V. rotiferianus* loads decreased rapidly over the same period and were undetectable (< 10^4^ copies/ng of DNA) in most individuals after 48 h (Fig. 3B). Second, we analyzed the effect of *Vibrio* strains on OsHV-1 growth. Remarkably, the viral load was 100-fold higher at 48h in oysters co-infected with *V. harveyi* and OsHV-1 than in oysters infected with OsHV-1 only (t-test, *p* < 0.05). Such an increase in viral load was not observed in co-infections with *V. rotiferianus* (Fig. 3C), consistent with the rapid elimination of *V. rotiferianus* from host tissues (Fig. 3D). Altogether, these results indicate that *V. harveyi* and OsHV-1 cooperate by increasing mutual growth during pathogenesis, an effect not observed with *V. rotiferianus*.

**Figure 3.**
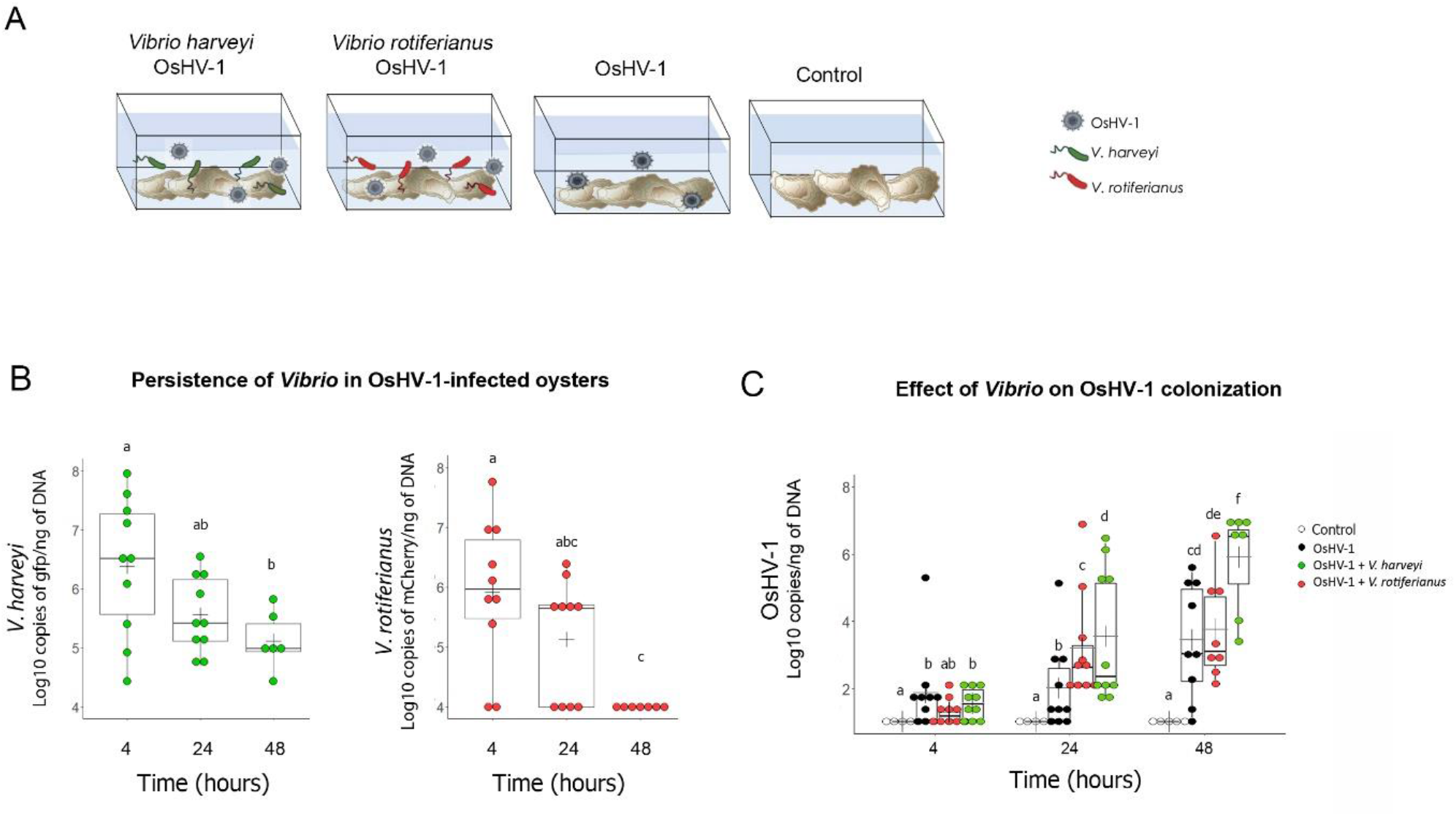
OsHV-1 and *V. harveyi* cooperate to colonize oysters. **(A) Simplified mesocosm experiment using fluorescent strains of *V. harveyi* and *V. rotiferianus* (Design 2)**. In order to identify putative cooperative behaviors between OsHV-1 and *Vibrio*, oysters were immersed at 20°C in seawater containing OsHV-1 and fluorescent *Vibrio* (either *gfp-*labeled *V. harveyi* Th15_O_G11 or *mCherry-*labeled *V. rotiferianus* Th15_O_G05). Exposure to OsHV-1 only or immersion in seawater without introduced pathogen were used as a control. Oysters were collected at 4 h, 24 h, and 48 h, *i*.*e*. before mortalities occurred, to monitor pathogen load in oyster tissues. **(B) Higher persistence of *V. harveyi* in OsHV-1-infected oysters**. *Vibrio* loads were determined by qPCR by quantifying *gfp* and *mCherry* copies in total DNA extracted from oyster flesh in OsHV-1/*V. harveyi* and OsHV-1/*V. rotiferianus* co-infections, respectively. Each dot represents an individual oyster (t-test, p < 0.05). Only *V. harveyi* is detected at > 10^5^ copies/ng DNA over the time course of the experiment. Detection limit: 10^4^ copies/ng DNA. **(C) *V. harveyi* promotes OsHV-1 replication**. OsHV-1 load was measured by qPCR in the flesh of oysters exposed to OsHV-1 and *gfp*-labeled *V. harveyi* Th15_O_G11 (green) or OsHV-1 and *mCherry-*labeled *V. rotiferianus* Th15_O_G05 (red), or OsHV-1 only (black). Each dot represents an individual. (t-test, p < 0.05). A significant increase in OsHV-1 load is observed in the presence of *V. harveyi*.

### *V. harveyi* actively dampens oyster immune defenses

We next investigated *Vibrio* traits that may facilitate host colonization and favor polymicrobial synergy with OsHV-1. We first compared the virulence potential of *V. harveyi* and *V. rotiferianus* (successful colonizers, Fig. 1B, 2C) and *V. jasicida* and *V. owensii* (poor colonizers). Virulence potential was tested *in vivo* by direct injection of *Vibrio* isolates into oyster adductor muscle. *V. harveyi* isolates showed strong virulence potential as revealed by an average of 50% oyster mortality one day after injection (Fig. 4A). However, mortalities remained below 15% on average after injection of *V. rotiferianus, V. jasicida*, and *V. owensii* (Fig. 4A). Not only did *V. harveyi* have a significantly higher virulence potential than other species (Kruskal-Wallis test, *p* < 0.001), but it also showed greater cytotoxicity toward oyster immune cells (hemocytes) *in vitro* (Fig. 4B). Here we compared the cytotoxic activity of two strains per *Vibrio* species among the four Harveyi-related species from our study. Upon exposure to *V. harveyi* strains, 67-76% of hemocytes underwent lysis (*p* < 0.001, One-way ANOVA, and Tukey post-hoc test) (Fig. 4B). The three other *Vibrio* species were not cytotoxic, with the percentage of lysed cells similar to controls (7 to 23%). We further confirmed the ability of *V. harveyi*, but not *V. rotiferianus*, to damage hemocytes using fluorescent *Vibrio* strains. After a 2 h exposure to *V. harveyi*, hemocytes were massively damaged and many extracellular *V. harveyi* were observed (Fig. 4C). In contrast, no cellular damage was observed when hemocytes were exposed to *V. rotiferianus*, with most bacteria being phagocytized. Thus, *V. harveyi* strains are equipped with specific virulence/cytotoxicity mechanisms that can dampen oyster cellular defenses. To uncover genomic features that might explain *V. harveyi* virulence/cytotoxicity, we sequenced and analyzed 17 Harveyi-related genomes from our collection (4 *V. harveyi*, 5 *V. owensii*, 4 *V. jasicida* and 4 *V. rotiferianus* isolates; Table S6). We found that the *V. harveyi* genome contained the most candidate virulence factors of the four *Vibrio* species in the present study. Candidate virulence genes in the *V. harveyi* genome included a type 3 secretion system (T3SS) and its associated effectors as well as 3-4 different type 6 secretion systems (T6SS) (Table S7). Such T6SS were previously shown to be essential for oyster colonization in *V. crassostreae* and *V. tasmaniensis* [19, 21].

**Figure 4.**
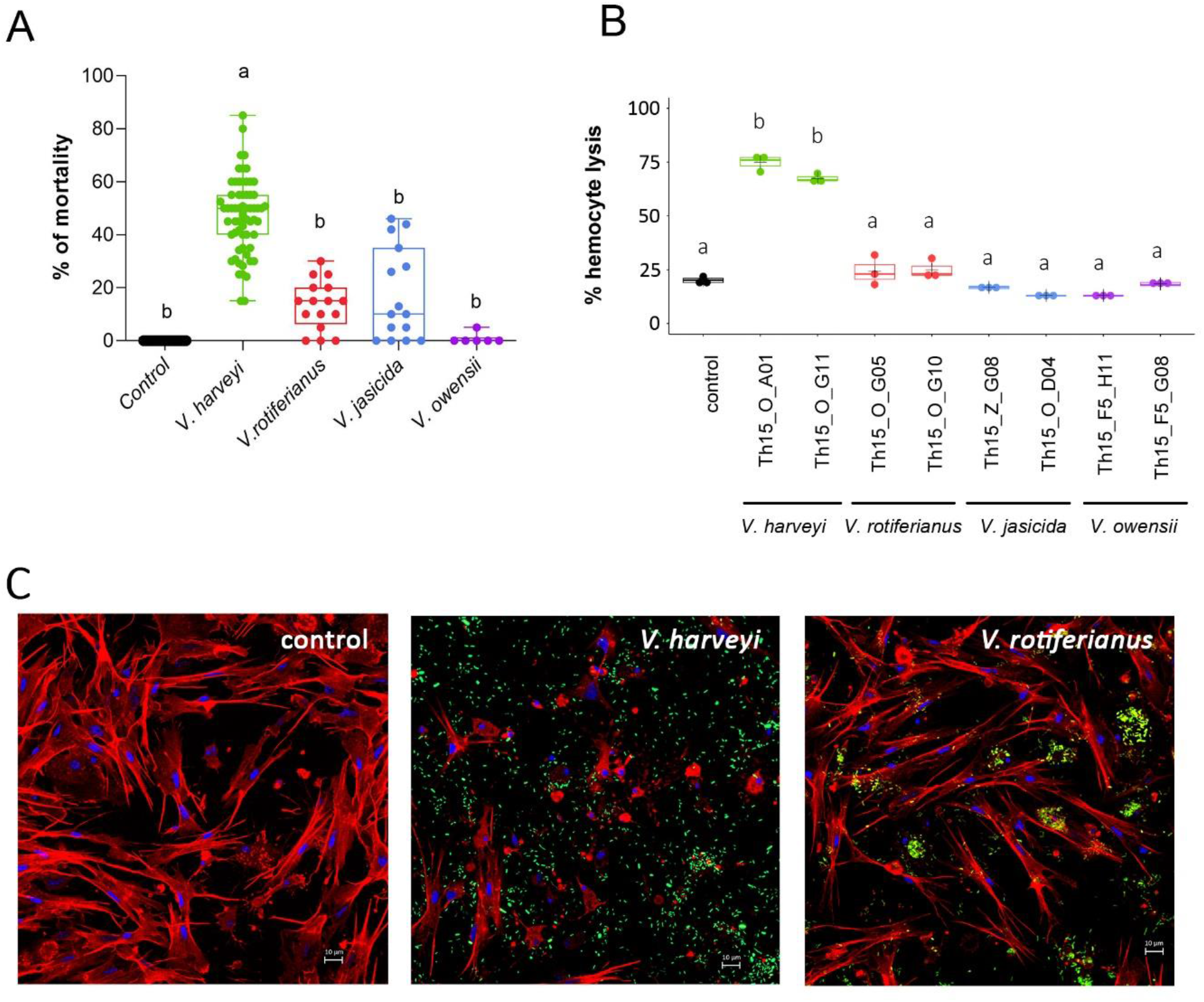
*V. harveyi* strains show high virulence potential and cytotoxicity toward oyster immune cells. **(A) *V. harveyi* is significantly more virulent than other Harveyi-related species in oyster experimental infection**. Oyster mortality rate (%) was measured at 24 h following an injection of Harveyi-related isolates. Mortality rates were compared for each *Vibrio* species using the Kruskal-Wallis test with post-hoc Dunn Test. Significant differences between mean values are represented by different letters (p-value < 0.0001). *V. harveyi* (n=63), *V. rotiferianus* (n=16), *V. jasicida* (n=15), *V. owensii* (n=6) (Kruskal-Wallis test, p < 0.001). *V. harveyi* isolates induced significantly higher mortalities than other strains. **(B) *V. harveyi* is significantly more cytotoxic to oyster hemocytes than other species**. *Vibrio* cytotoxicity was determined on monolayers of hemocytes and monitored using Sytox green labelling. Cells were incubated with bacteria at a MOI of 50:1. The maximum percentage of hemocyte lysis (%) caused by *Vibrio* is displayed. Error bars represent the standard deviation of the mean, different letters represent significant differences between means in a multiple comparison test (p<0.05, One-way ANOVA with post-hoc Tukey HSD Test). Strains of *V. harveyi* were significantly more cytotoxic than other strains. **(C) *V. harveyi* but not *V. rotiferianus* causes damage to oyster hemocytes**. The *Vibrio* effect on hemocytes was observed by epifluorescence microscopy. Monolayers of hemocytes were incubated with fluorescently-labeled *V. harveyi* Th15_O_G11 or *V. rotiferianus* Th15_O_G05 at a MOI of 50:1 for 2h. Actin was stained with Fluorescent-phalloidin (Red), Chromatin was stained with DAPI. *Vibrio* strains expressing fluorescent proteins are shown in Green. Cell damage was only observed with *V. harveyi*.

### *V. harveyi* produces vibrioferrin, which can be used by *V. rotiferianus*

We next used comparative genomics to examine how *V. harveyi* and *V. rotiferianus* both colonize OsHV-1-infected oysters but make distinct contributions to pathogenesis (Table S6). Using genome comparisons, 15 genes were identified that are exclusively shared by *V. harveyi* and *V. rotiferianus* (good colonizers; Table S8). Of these, iron acquisition systems differed between species (Fig. S7). We paid particular attention to genes involved in the vibrioferrin pathway, a siderophore whose uptake system was only found in strains of *V. harveyi* and *V. rotiferianus* (Fig. 5A). Vibrioferrin is a tricarboxylic acid siderophore derived from citric acid; it was shown to be shared as a public good within populations of *V. splendidus* [51]. Remarkably, we found that *V. harveyi* harbors genes responsible for the biosynthesis and uptake of vibrioferrin whereas *V. rotiferianus* only carries the vibrioferrin uptake system (Fig. 5A). This suggests that by secreting vibrioferrin, *V. harveyi* could facilitate iron uptake by *V. rotiferianus*. To determine whether *V. rotiferianus* is able to use vibrioferrin produced by *V. harveyi*, we compared the growth of *V. rotiferianus* in the presence/absence of increasing amounts of *V. harveyi* culture supernatant. *V. harveyi* was able to grow in iron-depleted medium (2,2’-bipyridine used as iron chelator, Fig. S8), in agreement with its ability to produce siderophores. In contrast, growth of *V. rotiferianus* was impaired upon iron depletion, (*p* < 0.05, Mann-Whitney) (Fig. S8). Growth of *V. rotiferianus* was rescued in a dose-dependent manner by the addition of cell-free culture supernatant from *V. harveyi* (Kruskal-Wallis, *p* < 0.05) (Fig. 5B). To determine whether this rescue is linked to vibrioferrin, we synthesized the siderophore (Fig. S9-S10). Synthetic vibrioferrin was sufficient to rescue *V. rotiferianus* growth in iron-depleted medium (Kruskal-Wallis, *p* < 0.05) (Fig. 5C). Finally, we constructed a *V. rotiferianus* mutant strain which lacks the two genes encoding the PvuA1-A2 receptor that are required for vibrioferrin uptake. Vibrioferrin failed to rescue the growth of the *V. rotiferianus* Δ*pvuA1-A2* mutant (Fig. 5D). This demonstrates that *V. rotiferianus* is able to acquire vibrioferrin, an important resource produced by *V. harveyi* (and potentially other *Vibrio* within the microbiota), to grow in iron-poor environments.

**Figure 5.**
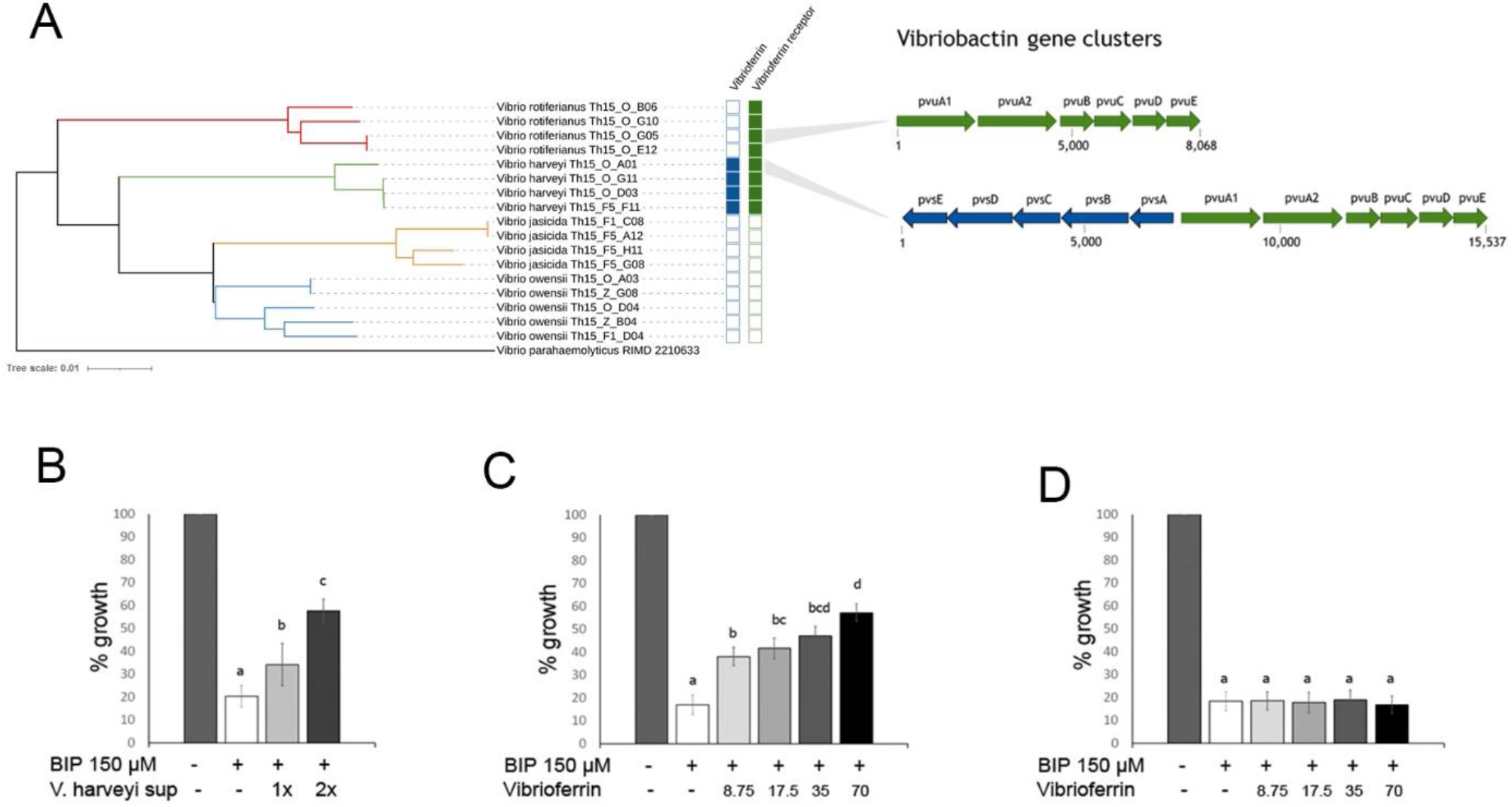
*V. rotiferianus* growth in iron-poor conditions is rescued by *V. harveyi* supernatant or vibrioferrin. **(A) Possible cheating on vibrioferrin uptake in *V. rotiferianus***. Comparative genomics between strains of *V. harveyi* (Th15_O_G11, Th15_F5_F11, Th15_O_D03, Th15_O_A01) and *V. rotiferianus* (Th15_O_G05, Th15_O_G10, Th15_O_B06, Th15_O_E12) point to a possible cheating behavior for vibrioferrin uptake in *V. rotiferianus*, which has receptor for vibrioferrin but does not produce it. **(B) *V. rotiferianus* growth in iron-depleted medium is rescued by *V. harveyi* culture supernatant**. Iron-depletion was obtained by adding 150 μM 2,2’-bipyridine (BIP) to the minimal culture medium. Dose-dependent growth rescue of strain Th15_O_G05 was achieved by adding *V. harveyi* Th15_O_G11 culture supernatant (either 1X or 2X concentrated by lyophilization). **(C) *V. rotiferianus* growth in iron-depleted medium is rescued by vibrioferrin**. Dose-dependent growth rescue of strain Th15_O_G05 was achieved by 8.75 to 70 μM vibrioferrin (see Fig. S8 for vibrioferrin synthesis). **(D) *V. rotiferianus* growth rescue requires the iron-vibrioferrin receptor PvuA**. Growth of *V. rotiferianus* Th15_O_G05 Δ*pvuA1-A2* in iron-depleted medium was not rescued by addition of vibrioferrin up to 70 μM. Letters indicate significant differences between conditions (p < 0.05, Kruskal-Wallis).

Overall, our data show that *V. rotiferianus* behaves as a cheater by using a siderophore produced by *V. harveyi*. Thus unlike *V. harveyi* and the OsHV-1 virus, which behave cooperatively, *V. rotiferianus* successfully colonizes oysters taking advantage of public goods contributed by the microbiota without providing benefit to the POMS-associated microbiota.

## Discussion

Here, we used a natural pathosystem, the Pacific Oyster Mortality Syndrome (POMS), to disentangle the complex web of interactions that shape polymicrobial assemblages and pathogenicity. Our data highlighted a web of interdependencies in which diverse microorganisms shape the POMS pathobiota and accelerate disease progression. In particular, we identified conserved cooperative traits accelerating disease progression as well as incidental cheating traits. These traits contribute to shaping the *Vibrio* community associated with OsHV-1-infected oysters.

We showed here that the population structure of *Vibrio*, a genus consistently found in the POMS pathobiota [52], varies across French Atlantic and Mediterranean oyster-farming environments. Indeed, we found *V. harveyi* and *V. rotiferianus* (Harveyi clade) positively associate with OsHV-1-infected oysters during field mortalities in the Mediterranean. This finding contrasts with a previous analysis of Atlantic oysters where *V. crassostreae* (Splendidus clade) dominated [17]. Experimentally, two out of four different species from the same clade were recruited from seawater, suggesting that the environment serves as a reservoir for *Vibrio* recruitment during POMS. Differences in *Vibrio* species associated with OsHV-1-infected oysters are therefore likely related to different environmental distributions of *Vibrio* species. Indeed, while *V. crassostreae* is present at high latitudes, e.g. in the North sea, Germany [53], *V. harveyi* preferentially grows in warm waters [54], such as those found for half the year in the Thau lagoon (16-30°C) [55]. Consistently, *V. harveyi* has been associated with disease outbreaks under rising sea surface temperatures [56]. Remarkably, the data from both locations (Atlantic/Mediterranean) converges in that the OsHV-1 virus is associated with specific populations of *Vibrio* in diseased oysters only. Our data indicate that OsHV-1 is key to enabling colonization by such specific *Vibrio* species.

By focusing on *Vibrio* that naturally co-infect oysters with OsHV-1, we observed a first level of interdependence and polymicrobial synergy occurring during POMS. Indeed, the combined effects of OsHV-1 and *Vibrio* triggered a faster host death than that observed when the microorganisms were used in isolation to infect oysters. While both *V. harveyi* and *V. rotiferianus* successfully colonized OsHV-1-infected oysters, polymicrobial synergy was specifically observed between OsHV-1 and *V. harveyi*. The benefits of co-infection were not only observed for OsHV-1 but also for the whole *Vibrio* community: both showed significantly greater expansion prior to oyster death when OsHV-1 and *V. harveyi* were co-infected. Whether other bacterial genera conserved among POMS consortia (*e*.*g. Arcobacter, Marinomonas*) [15, 52, 57] also contribute to this polymicrobial synergy or play complementary roles in the polymicrobial consortium remains to be established.

A key mechanism underlying polymicrobial synergy between OsHV-1 and *Vibrio* is the dampening of oyster cellular defenses, which makes the local environment less hostile for the entire microbiota. This manipulation of cellular immunity is key in a number of polymicrobial infections. For instance, induction of IL-10-producing macrophages by *M. tuberculosis* favors HIV-1 replication and spread [8]. Similarly, the apoptosis of macrophages, neutrophils, dendritic cells and NK cells contributes to the fatal outcome of influenza A virus / *S. pneumoniae* co-infections [5]. Importantly, cytotoxicity to host immune cells is a phenotypic trait conserved in *Vibrio* species that co-occurs with OsHV-1 in diseased oysters. Indeed, cytotoxic effects toward oyster hemocytes is a functional trait conserved in Mediterranean strains of *V. harveyi*, as well as in strains of *V. crassostreae, V. tasmaniensis* and *V. splendidus* [19, 21, 58] isolated from POMS-diseased oysters in the Atlantic [17, 20, 25, 59, 60]. We previously showed that cytotoxicity toward immune cells is essential for *Vibrio* to colonize oyster tissues [21]. This indicates that *Vibrio* species that harbor similar functions are likely replaceable within the POMS bacterial consortia. Our mesocosm experiment validated the preferential association of *V. harveyi* with OsHV-1-infected oysters under controlled conditions, showing that cytotoxic *Vibrio* species can be recruited from the ecosystems where they circulate. In the mutualistic association between the mollusk *Euprymna scolopes* and its symbiont *Vibrio fischeri*, it was also shown that *Vibrio* phenotypic traits determine their capacity to be recruited from the environment [61]. Overall, our data show that similar to OsHV-1 [14], cytotoxic *Vibrio* species modify their extracellular environment by targeting oyster immunity. This results in eased proliferation of co-infecting partners, as shown here for OsHV-1 and specific *Vibrio* species. This fundamental cooperation taking place within the POMS pathobiota has benefits for the entire polymicrobial consortium.

Our present study suggests that cooperation through dampening host defenses contributes to the shaping of the POMS pathobiota assembly. This supports recent studies highlighting that cooperative interactions within animal microbiota participate in the shaping of community composition and functioning (for review see [62]). However, pathogenic microbes cooperating to manipulate their host may be outcompeted by other members of the microbiota that do not invest in cooperation. Manipulation is therefore only expected to be favored if its benefits predominantly fall back on the manipulator [63]. One important benefit for OsHV-1 and *V. harveyi* observed here is a higher load for both manipulators. Another likely benefit is increased pathogen transmission due to accelerated host death (Fig. 6). From a general point of view, accelerating host death is not predicted to be favorable to pathogens as this may drive the host, and consequently themselves, to extinction, unless transmissibility is also increased [64]. This is particularly true for pathogens with narrow host spectra like *V. harveyi* and *V. crassostreae*, but also for OsHV-1, which are almost exclusively found in oysters (this study;[17]). POMS is typically transmitted from oyster to oyster by a massive release of pathogens into the seawater, which in turn infects neighboring oysters through filter-feeding [57, 65]. We can therefore hypothesize that polymicrobial synergy leading to accelerated release of OsHV-1 and *Vibrio* into seawater is advantageous in terms of group selection.

**Figure 6.**
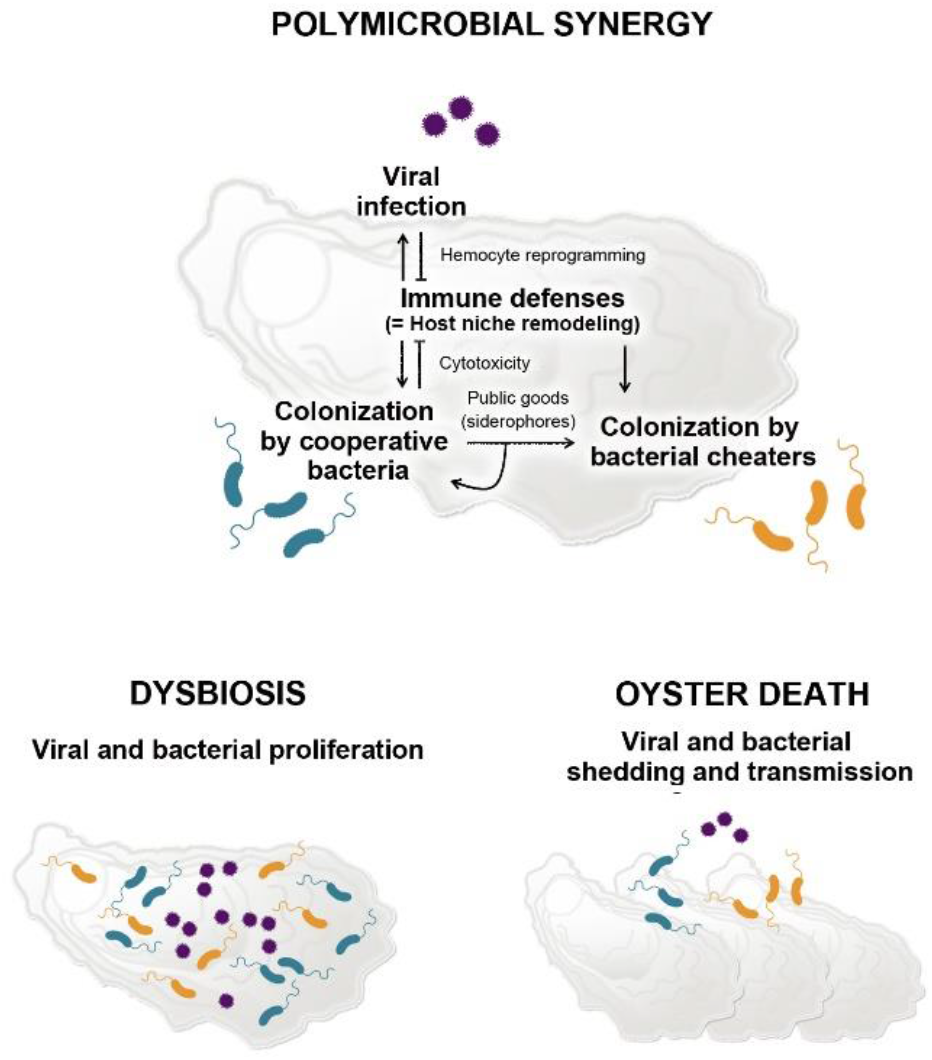
A systemic view of Pacific Oyster Mortality Syndrome polymicrobial synergy. Oysters are infected by OsHV-1 μvar, which impairs host immune defenses making the environment less hostile for stable bacterial colonization. Secondary bacterial colonization is enabled. The subset of cooperative bacteria that exhibit cytotoxicity toward hemocytes further dampen oyster immune defenses. This dampening is beneficial to the whole microbial community (bacteria and the OsHV-1 virus) and leads to dysbiosis. Cooperative bacteria also secrete siderophores that can be used by other members of the community who do not invest in costly mechanisms of colonization (cheaters). Cooperation between the virus and cytotoxic bacteria accelerates host death leading to the shedding of bacteria and viruses that can infect new hosts.

Beyond cooperative traits conserved among successful *Vibrio* colonizers, we find evidence that cheating is an efficient strategy that *Vibrio* use to colonize oysters affected by POMS. Indeed, unlike *V. harveyi, V. rotiferianus* does not invest resources in virulence nor cytotoxicity. Moreover, our data argues that *V. rotiferianus* lacks the costly pathways to synthesize vibrioferrin. Instead, this species imports this siderophore, produced by *V. harveyi*, to promote its own growth. In contrast, the unsuccessful colonizers *V. owensii* and *V. jasicida* lack the vibrioferrin uptake machinery, which may contribute to their competitive exclusion [66].

Remarkably, vibrioferrin biosynthetic pathways are highly conserved between *V. harveyi* and *V. crassostreae*, the two main species associated with POMS in the Mediterranean and the Atlantic. Comparison of these *Vibrio* genomes reveals conserved synteny and > 70% amino acid identity in homologous proteins involved in these pathways (Fig. S12). Both *Vibrio* species can, therefore, supply similar public goods to the POMS pathobiota. Metabolite cross-feeding, which enables a bacterium to consume metabolites produced by another community member, can mediate synergy in multi-species infections (for review see [1]). Cheating behavior regarding iron-acquisition was earlier described in *Vibrio* by Cordero *et al*. [51]. The authors showed that within ecologically cohesive clusters of closely related *Vibrio*, only some genotypes were able to produce siderophores. Meanwhile, non-producers had selectively lost siderophore biosynthetic pathways. We observe traces of selective loss in *V. rotiferianus* strains: while *V. rotiferianus* and *V. harveyi* both harbor the genomic region that contains vibrioferrin receptors, the former species specifically lacks the vibrioferrin biosynthetic genes.

The cheating behavior arguably coevolves with ecological adaptation of *Vibrio* toward association with larger particles in the water column, consistent with efficient siderophore sharing where local densities of bacteria are high [51]. Because oysters host dense populations of bacteria in their body fluids (> 10^7^ culturable bacteria/mL during episodes of POMS [67]), they constitute microhabitats where cheating can occur. Oysters could, therefore, provide a favorable niche for *Vibrio* strains that can import iron-loaded siderophores. Importantly, the present study shows that cheating for iron acquisition occurs inter-specifically and coincides with the co-occurrence of *V. harveyi* and *V. rotiferianus* in POMS-diseased oyster. Indeed, our two-year Mediterranean field survey showed that *V. harveyi* is repeatedly associated with diseased oysters. *V. harveyi* co-occurs with *V. rotiferianus* in oyster flesh, and to a lower extent in particles >60μm (zooplankton) and 5-60μm (phytoplankton). These host-associated states provide conditions where *V. rotiferianus* could have evolved cheating behavior. Thus, social interactions through siderophore-sharing appear to structure the assembly of bacterial species in oysters affected by POMS.

## Conclusion

By disentangling the complex interactions at play in the Pacific Oyster Mortality Syndrome, we have shown that cooperation is key in the functioning of this natural pathosystem. Cooperation and cheating seem to drive the assembly of the pathobiota. The former manifests as polymicrobial synergy between the OsHV-1 virus, the etiological agent of POMS and secondary *Vibrio* colonizers. Dampening of oyster cellular defenses and siderophore sharing, both of which make the host environment more favorable for microbial proliferation, are two cooperative traits conserved in the main POMS-associated *Vibrio* species, namely *V. harveyi* and *V. crassostreae*, across oyster farming environments. This knowledge opens new avenues for the control of polymicrobial diseases by interfering with polymicrobial assembly. Implementation of ecological principles, such as interfering with cooperative behavior within the microbiome (*e*.*g*. siderophore sharing) or altering the local environment of the POMS pathobiota (*i*.*e*. stimulating host defenses) are promising solutions for future exploration. We have shown recently that eliciting oyster antiviral defenses through so-called immune priming is sufficient to prevent colonization by OsHV-1 and subsequent disease development [37, 68]. Similarly, exposure to microbial communities at early developmental stages (referred to as biological embedding) was also protective against POMS [69]. Such solutions, which make the host a less favorable niche for the OsHV-1 virus [68] and/or bacteria [69], are promising avenues for preventing or reducing the assembly of microbial consortia that produce POMS.

## Declarations

### Ethics approval

The animal (oyster *Crassostrea gigas*) testing followed all regulations concerning animal experimentation. The authors declare that the use of genetic resources fulfill the French regulatory control of access and EU regulations on the Nagoya Protocol on Access and Benefit-Sharing (ABSCH-IRCC-FR-259502-1).

### Availability of data and materials

Targeted gene sequences (*hsp60, rctB, topA* and *mreB)* and all sequence files with associated metadata generated in mesocosm experiments are available in Ifremer Oceanic database (doi doi.org/10.12770/173c0414-a3ca-4a79-b6b2-cd424ee90593 [28] and doi.org/10.12770/63b02659-cd9d-4834-8e6d-8adfa736755d [70]). Genome assemblies have been deposited in the European Nucleotide Archive (ENA) under project accession no. PRJEB49488. Original R statistic scripts for metagenomics analyses and the phyloseq table are available https://doi.org/10.5281/zenodo.7599486.

### Author contributions

Conception: DDG, MAT, GMC, FLR, GM. Designed the work: DDG, MAT, GMC, FLR, GM, DO, AL. Acquisition, analysis and/or interpretation of data: DO, AL, MB, PH, AM, YD, FD, NI, BM, ET, CC, CM, JME, YL, YG, JDL, LD, BP, DT, LLP, ML, OR, JP, GM, FLR, GMC, MAT, DDG. Drafted the work: DDG, MAT, GMC, DO, AL or substantively revised it: FLR, GM, JME JDL.

## Acknowledgments

We thank Jean-François Allienne at the Bioenvironment platform (University Perpignan Via Domitia) for supporting NGS library preparation and sequencing as well as Valentin Outrebon and Anna Amouroux for crucial technical help. We thank Montpellier RIO Imaging (https://www.mri.cnrs.fr) for access to microscopy facilities and the qPHD platform/Montpellier genomix for access to qPCR. We thank the Regional Committee of Mediterranean Shellfish Aquaculture (CRCM) and the Ifremer for access to the shellfish tables and for boats. We thank the GENSEQ platform (http://www.labex-cemeb.org/fr/genomique-environnementale-2) from the labEx CeMEB for nucleotide sequencing.

