## Supplementary material for "Cooperation and cheating orchestrate *Vibrio* assemblages and polymicrobial synergy in oysters infected with OsHV-1 virus"

**Table S1. Prevalence of the Harveyi clade in oyster flesh during Mediterranean POMS episodes.**

Oysters were collected during and outside POMS episodes in the Thau lagoon. At each sampling time, a total of 76 to 96 bacterial isolates obtained from oyster flesh were screened for the Harveyi clade (rctB-F/rctB-R primers) and the *V. harveyi* species (Vh-mreB-F/Vh-mreB-R primers). The oyster flesh was simultaneously tested for OsHV-1 µvar detection (OsHVDPFor/OsHVDPRev primers).

| Date | Field mortalities | OsHV-1 µvar virus | % Harveyi clade | % *V. harveyi species* |
|---|---|---|---|---|
| **1-Oct-2015** | Yes | positive | 94.0 | 75.9 |
| **9-June-2016** | No | negative | 0.0 | 0.0 |
| **23-June-2016** | Yes | positive | 49.0 | 32.0 |
| **13-March-2017** | No | negative | 1.0 | 1.0 |

**Table S2. Samplings in the Thau lagoon.**

<table>
<tr><th>Sampling date</th><th>GPS coordinates</th><th>Temp</th><th>Juvenile Mortality</th><th>Biological material</th><th>Hachery</th><th>Time of immersion</th><th colspan="2">No of isolates</th><th colspan="2">No of Vibrio isolates*</th><th colspan="2">No of confirmed Harveyi isolates**</th></tr>
<tr><td>1-Oct-2015</td><td>43°25'07.5"N 3°37'21.4"E</td><td>18°C</td><td>yes</td><td>Oysters (n=30)</td><td>Ifremer<br>France naissain</td><td>4 weeks</td><td>96</td><td rowspan="5">472</td><td>55</td><td rowspan="5">437</td><td>54</td><td></td></tr>
<tr><td>1-Oct-2015</td><td>43°25'07.5"N 3°37'21.4"E</td><td>18°C</td><td>yes</td><td>Water column > 60µm</td><td>n/a</td><td>n/a</td><td>96</td><td>59</td><td>5</td><td rowspan="4">53</td></tr>
<tr><td>1-Oct-2015</td><td>43°25'07.5"N 3°37'21.4"E</td><td>18°C</td><td>yes</td><td>Water column 5-60 µm</td><td>n/a</td><td>n/a</td><td>96</td><td>61</td><td>21</td></tr>
<tr><td>1-Oct-2015</td><td>43°25'07.5"N 3°37'21.4"E</td><td>18°C</td><td>yes</td><td>Water column 1-5 µm</td><td>n/a</td><td>n/a</td><td>96</td><td>67</td><td>23</td></tr>
<tr><td>1-Oct-2015</td><td>43°25'07.5"N 3°37'21.4"E</td><td>18°C</td><td>yes</td><td>Water column 0,2-1 µm</td><td>n/a</td><td>n/a</td><td>88</td><td>62</td><td>4</td></tr>
<tr><td>9-June-2016</td><td>43°22'44.8"N 3°34'17.5"E</td><td>24°C</td><td>no</td><td>Oysters (n=30)</td><td>Ifremer</td><td>1 week</td><td>76</td><td></td><td>76</td><td></td><td>0</td><td></td></tr>
<tr><td>23-June-2016</td><td>43°22'45.1"N 3°34'16.1"E</td><td>22°C</td><td>yes</td><td>Oysters (n=29)</td><td>Ifremer</td><td>10.5 weeks</td><td>96</td><td></td><td>96</td><td></td><td>47</td><td></td></tr>
<tr><td>13-March-2017</td><td>43°26.058'N 003°39.878'E</td><td>12.6°C</td><td>no</td><td>Oysters (n=12)</td><td>Ifremer</td><td>2 weeks</td><td>96</td><td></td><td>96</td><td></td><td>1</td><td></td></tr>
</table>

* based on *hsp60* sequencing

** based on MLST (*hsp60, rctB, topA*, *mreB)*

**Table S3. List of primers**

| Primer name | Primer sequence | Description and references |
|---|---|---|
| rctB-Fw-I | 5'-TCGTCGGCAGCGTCAGATGTGTATAAGA-GACAGYRTGAATAGGCTCAAATTCGCCGTC -3' | Forward primer for *rctB* gene amplification (barcoding), this study |
| rctB-Rv-I | 5'- GTCTCGTGGGCTCGGAGATGTGTATAAGAGA-CAGYRCCWTCSTTCTWYGAAGAAYYGCT -3' | Reverse primer for *rctB* gene amplification (barcoding) , this study |
| Insert-pLP12-R | 5'- CGGCTGACATGGGAATTGC -3' | Forward primer annealing upstream of the multiple cloning site of pLP12, this study |
| del-pvuA1-A2-OG05-F | 5'- GTCATGCTCACGACCAAAC -3' | Forward primer annealing upstream of *pvuA1.* Used to verify integration of pAM010 in *V. rotiferianus* Th15_O_G05 and to verify the presence of *pvuA1* and *pvuA2* genes, this study |
| del-pvuA1-A2-OG05-R | 5'- GGCTTGCTCACTTGGTTG -3' | Reverse primer annealing downstream of *pvuA2.* Used to verify the presence of p*vuA1* and *pvuA2* genes in *V. rotiferianus* Th_15_O_G05, this study |
| hsp60-F | 5'- GAATTCGAIIIIGCIGGIGAYGGIACIACIAC -3' | Forward primer for hsp60 gene amplification (MLST) [1] |
| hsp60-R | 5'- CGCGGGATCCYKIYKITCICCRAAICCIGGIGCYTT -3' | Reverse primer for hsp60 gene amplification (MLST), [1] |
| VtopA400F | 5'- GAGATCATCGGTGGTGATG -3' | Forward primer for topA gene amplification (MLST) [2] |
| VtopA1200R | 5'- GAAGGACGAATCGCTTCGTG -3' | Reverse primer for topA gene amplification (MLST) [2] |
| rctB-F | 5'- GCTGATGAAAAAATTCTGATTAAAGC -3' | Forward primer for Harveyi clade rctB gene amplification (MLST), this study |
| rctB-R | 5'- TGAATAGGCTCAAATTCGCCGTC -3' | Reverse primer for Harveyi clade rctB gene amplification (MLST), this study |
| VmreB12F | 5'- ACTTCGTGGCATGTTTTC -3' | Forward primer for mreB gene amplification (MLST) [2] |
| VmreB999R | 5'- CCGTGCATATCGATCATTTC -3' | Reverse primer for mreB gene amplification (MLST) [2] |
| OsHVDPFor | 5'- ATTGATGATGTGGATAATCTGTG -3' | Forward primer for OsHV-1 DNA pol gene amplification (quantification) [3] |
| OsHVDPRev | 5'- GGTAAATACCATTGGTCTTGTTCC -3' | Reverse primer for OsHV-1 DNA pol gene amplification (quantification) [3] |
| Vh-mreB-F | 5'- GCTGATGAAAAAATTCTGATTAAAGC -3' | Forward primer for *V. harveyi* mreB gene amplification (detection), this study |

| | | |
|---|---|---|
| Vh-mreB-R | 5'- TGAATAGGCTCAAATTCGCCGTC -3' | Reverse primer for *V. harveyi* mreB gene amplification (detection), this study |
| 567F | 5'- GGCGTAAAGCGCATGCAGGT -3' | Forward primer for Vibrio 16s rDNA amplification (quantification) [4] |
| 680R | 5'- GAAATTCTACCCCCCTCTACAG -3' | Reverse primer for Vibrio 16s rDNA amplification (quantification) [4] |
| GFP_F522 | 5'- TGGAAGCGTTCAACTAGCAG -3' | Forward primer for *gfp* amplification (quantification), This study |
| GFP_R625 | 5'- AAAGGGCAGATTGTGTGGAC -3' | Reverse primer for *gfp* amplification (quantification), This study |
| mCherry_F494 | 5'- AGATCAAGCAGAGGCTGAAGCTGA -3' | Forward primer for *mCherry* amplification (quantification) [5] |
| mCherry-R655 | 5'- ACTGTTCCACGATGGTGTAGTCCT -3' | Forward primer for *mCherry* amplification (quantification) [5] |
| hr-F1 | 5'- AAGAGGTTCAAGCCCAGACG -3' | Forward primer for *V. harveyi-V. rotiferianus* chemotaxis protein amplification (quantification), This study |
| hr-R1 | 5'- AGTCCTGCGCTTGTTTCTCA -3' | Forward primer for *V. harveyi-V. rotiferianus* chemotaxis protein amplification (quantification), This study |
| oj-F1 | 5'- CCAGAAGAGCAAACCGTGAT -3' | Forward primer for *V. owensii-V. jasicida ompA* amplification (quantification), This study |
| oj-R1 | 5'- CGACTGGCTAACTGCATCAA -3' | Reverse primer for *V. owensii-V. jasicida ompA* amplification (quantification), This study |

**Table S4. List of genetically modified strains**

| Strains | Description | Reference |
|---|---|---|
| ***E. coli*** | | |
| DH5αλpir | *sup* E44, Δ*lacU*169 (Φ*lacZ*ΔM15), *recA1*, *endA1*, *hsdR17*, *thi-1*, *gyrA96*, *relA1*, λpir phage lysogen | LEMAR collection |
| β3914 | F-, RP4-2-Tc::Mu, Δ*dapA*::(*erm*-pir) *gyrA462* zei-298::Tn10, $Km^R$, $Em^R$, $Tc^R$, DAP- | [6] |
| GEB883 + pEVS104 | Strain GEB883 containing pEVS104. Used as a helper strain during bacterial conjugation | [7] |
| DH5α λpir + pAM010 | *E. coli* DH5α λpir containing the pAM010 | This study |
| β3914 + pAM010 | *E. coli* β3914 containing the pAM010 | This study |
| DH5α λpir + pFD086 | *E. coli* DH5α λpir containing the pFD086 (GFP, $Trim^R$) | [7] |
| DH5α λpir + pFD096 | *E. coli* DH5α λpir containing the pFD096 (mCherry, $Trim^R$) | This study |
| ***V. rotiferianus*** | | |
| Th15_O_G05 | *V. rotiferianus* strain, isolated in 2015 in oysters sampled in Thau lagoon | This study |
| Th15_O_G05 ΔpvuA1-2 | *V. rotiferianus* O_G05 strain in which the *pvuA1* and the *pvuA2* genes have been deleted | This study |
| Th15_O_G05 mCherry | *V. rotiferianus* O_G05 strain in which the pFD096 plasmid was conjugated (mCherry, $Trim^R$) | This study |
| ***V. harveyi*** | | |
| Th15_0_G11 | *V. harveyi* strain, isolated in 2015 in oysters sampled in Thau lagoon | This study |
| Th15_O_G11 GFP | *V. harveyi* O_G11 strain in which the pFD086 plasmid was electroporated (GFP, $Trim^R$) | This study |

**Table S5. List of plasmids used for mutagenesis and labelling**

| Plasmid name | Description | Reference |
|---|---|---|
| pMK-RQ_updn-pvuA1-2 | Cloning vector in which the upstream region of *pvuA1* and the downstream region of *pvuA2* of *V. rotiferianus* Th15_O_G05 have been cloned. $Km^R$ | GeneArt |
| pLP12 | Suicide plasmid used for targeted deletion of chromosomal genes of *Vibrio* species. oriT-RP4, oriV-R6K, $P_{BAD}$-*vmi480*. $Cm^R$ | [8] |
| pAM010 | Derivative of pLP12 in which the upstream region of *pvuA1* and the downstream region of *pvuA2* of *V. rotiferianus* Th15_O_G05 have been inserted. $Cm^R$ | This study |
| pFD086 | Replicative plasmid for *Vibrio* species containing the *gfp* gene under the control of the constitutive $P_{lac}$ promoter. $Trim^R$ | [7] |
| pFD096 | Derivative plasmid of pFD086 in which the *gfp* gene has been replaced by the *mcherry* gene of pME9407 (Rochat *et al.*, 2010), using the XbaI and BamHI sites. In silico plasmid map is available upon request. | This study |
| pMRB | High copy number replicative plasmid. $P_{lac}$-GFP, $Cm^R$ | [9] |

**Table S6. Strains sequenced by high-throughput sequencing.**
In this study, 17 Harveyi-related genomes from our collection (4 *V. harveyi*, 5 *V. owensii*, 4 V*. jasicida* and 4 *V. rotiferianus* isolates) were sequenced. The genome sequences have been deposited in the European Nucleotide Archive (ENA) under project accession number PRJEB49488.

| Species | Isolates | Origin | Contig number | Genome size (Mb) | CDSs | Accession number |
|---|---|---|---|---|---|---|
| ***V. harveyi*** | Th15_O_G11 | This study | 54 | 5.88 | 5,485 | ERS9919775 |
| | Th15_F5_F11 | This study | 106 | 5.96 | 5,615 | ERS9919777 |
| | Th15_O_D03 | This study | 64 | 5.50 | 5,497 | ERS9919776 |
| | Th15_O_A01 | This study | 99 | 5.89 | 5,507 | ERS9780105 |
| ***V. rotiferianus*** | Th15_O_G05 | This study | 85 | 5.29 | 4,968 | ERS9919778 |
| | Th15_O_G10 | This study | 81 | 5.16 | 4,862 | ERS9919779 |
| | Th15_O_B06 | This study | 79 | 5.17 | 4,863 | ERS9919780 |
| | Th15_O_E12 | This study | 45 | 5.33 | 5,015 | ERS9919781 |
| ***V. jasicida*** | Th15_F5_H11 | This study | 52 | 6.1 | 5,724 | ERS9919787 |
| | Th15_F1_A12 | This study | 159 | 7.12 | 7,262 | ERS9919788 |
| | Th15_F1_C08 | This study | 158 | 7.12 | 7,251 | ERS9919789 |
| | Th15_F5_G08 | This study | 61 | 5.89 | 5,431 | ERS9919790 |
| ***V. owensii*** | Th15_Z_G08 | This study | 98 | 6.02 | 5,578 | ERS9919782 |
| | Th15_O_D04 | This study | 46 | 5.95 | 5,473 | ERS9919783 |
| | Th15_O_A03 | This study | 95 | 6.01 | 5,568 | ERS9919784 |
| | Th15_Z_B04 | This study | 43 | 5.97 | 5,474 | ERS9919785 |
| | Th15_F1_D04 | This study | 75 | 6.25 | 5,870 | ERS9919786 |

Abbreviation; CDS, coding DNA sequence.

**Table S7. Virulence and candidate virulence genes found in Vibrio strains used in mesocosm experiments.**

X indicates the presence of a T3SS and/or a T6SS in isolates. Among the T3SS1 associated effectors characterized in Harveyi isolates, we identified VopQ, VopR, VopS and VPA0450 as well as an associated cargo chaperone orthologous to VPA0451, which is needed for translocation of VPA0450 into the host cell membrane [10].

| Species | Isolates | % mortality at day 1 | T3SS | T3SS effectors | T6SS1 | T6SS2 | T6SS3 | T6SS4 |
|---|---|---|---|---|---|---|---|---|
| ***V. harveyi*** | Th15_O_G11 | 50 | X | VopQ, VopR, VPA0450, VPA0451 | X | X | X | |
| | Th15_F5_F11 | 41 | X | VopQ, VopR, VPA0450, VPA0451 | X | X | X | X |
| | Th15_O_D03 | 28 | X | VopQ, VopR, VPA0450, VPA0451 | X | X | X | |
| | Th15_O_A01 | 46 | X | VopQ, VopR, VPA0450, VPA0451 | X | X | X | |
| ***V. rotiferianus*** | Th15_O_G05 | 30 | | | X | | X | X |
| | Th15_O_G10 | 10 | | | X | | X | X |
| | Th15_O_B06 | 0 | | | X | | X | X |
| | Th15_O_E12 | 15 | | | X | | X | X |
| ***V. jasicida*** | Th15_F5_H11 | 44 | X | VopQ, VopS | X | X | X | |
| | Th15_F1_A12 | 26 | X | VopQ, VopS | X | X | X | |
| | Th15_F1_C08 | 28 | X | VopQ, VopS | X | X | X | |
| | Th15_F5_G08 | 42 | X | VopQ, VopS | X | X | X | |
| ***V. owensii*** | Th15_Z_G08 | 0 | | | X | X | X | |
| | Th15_O_D04 | 0 | | | X | X | X | |
| | Th15_O_A03 | 0 | | | X | X | X | |
| | Th15_Z_B04 | 0 | | | X | X | X | |
| | Th15_F1_D04 | 0 | | | X | X | X | |

**Table S5. List of *V. harveyi* and *V. rotiferianus* specific genes.**
As example, genes corresponding to the isolate *V. harveyi* Th15_O_A1 are indicated in this list.
Orthologues of all these genes were identified only in *V. harveyi* and *V. rotiferianus* isolates.

| Vibrio harveyi Th15_O_A1 | Product | Function |
|---|---|---|
| VHARVA1S8_v1_100126 | protein of unknown function | |
| VHARVA1S8_v1_190050 | Chemotaxis protein | |
| VHARVA1S8_v1_260075 | protein of unknown function | |
| VHARVA1S8_v1_260092 | conserved membrane protein of unknown function | |
| VHARVA1S8_v1_360021 | conserved protein of unknown function | |
| VHARVA1S8_v1_460139 | conserved exported protein of unknown function | |
| VHARVA1S8_v1_480079 | conserved exported protein of unknown function | |
| VHARVA1S8_v1_480080 | Peptidase M66 family protein | |
| VHARVA1S8_v1_480410 | conserved protein of unknown function | |
| VHARVA1S8_v1_530104 | PvuA1, TonB-dependent siderophore receptor family protein | Vibrioferrin transport |
| VHARVA1S8_v1_530105 | PvuA2, TonB-dependent siderophore receptor family protein | |
| VHARVA1S8_v1_530106 | PvuB, iron-dicitrate transporter subunit ; periplasmic-binding component | |
| VHARVA1S8_v1_530285 | conserved membrane protein of unknown function | |
| VHARVA1S8_v1_80159 | conserved protein of unknown function | |
| VHARVA1S8_v1_80182 | glycosyltransferase family 25 protein | |

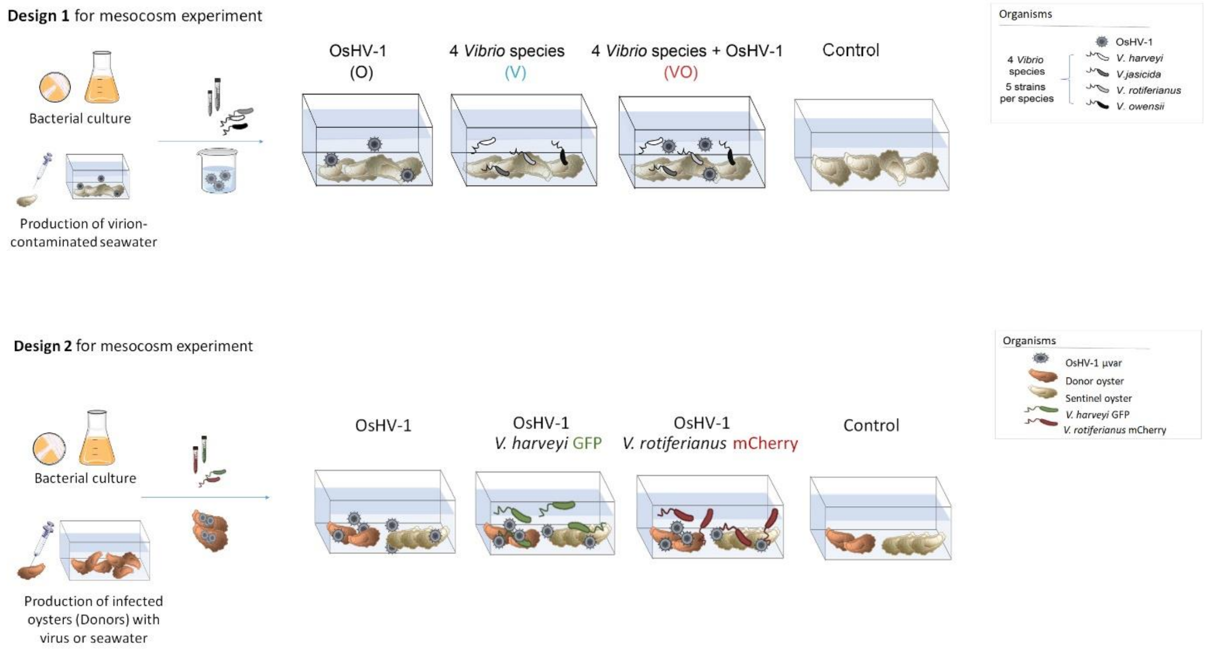

**Fig S1. Experimental designs for mesocosm experiments**

**Design of mesocosm experiments to mimic the polymicrobial disease. In design 1**, oysters were immersed at 20°C in contaminated seawater (1) with OsHV-1 virions freshly excreted from oysters ($2 \times 10^3$ copies/µl), (2) with 20 cultured vibrio (5 strains x 4 species of the Harveyi clade: *V. harveyi, V. rotiferianus, V. owensii, V. jasicida* at $10^7$ bacteria/ml), or (3) both. **In design 2**, oysters were immersed at 20°C firstly with OsHV1-contaminated source oysters. 24 hours later, GFP-tagged cultured *V. harveyi* or mCherry-tagged cultured *V. rotiferianus* were added in tanks to reach a final concentration of $10^7$ bacteria/ml. In both designs, oyster mortalities were monitored daily over 6 days. 10 individuals were sampled at each time point (0, 4, 24 and 48 h) for monitoring pathogens and the host response. Control tanks correspond to uninfected animals.

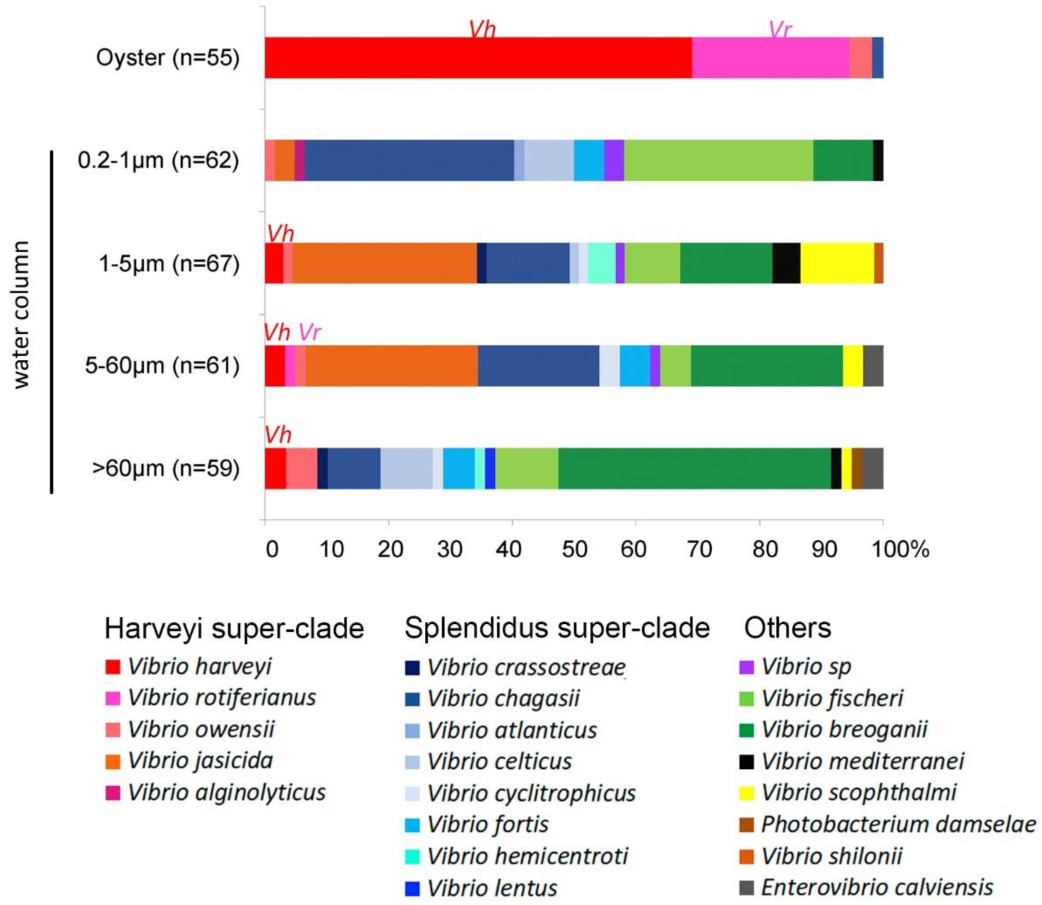

**Fig S2: *Vibrio* community structure in seawater fractions**

The population structure of *Vibrionaceae* was determined in seawater and oyster flesh during an episode of POMS. The graph shows the distribution (%) of *Vibrio* isolated from oysters and size-filtered seawater fractions (>60 µm, 60-5 µm, 5-1 µm, 1-0.2 µm), from which it stands out the high prevalence of the Harveyi super-clade (*V. harveyi* –*Vh* and *V. rotiferianus* –*Vr*) in oysters. Altogether, 304 *hsp60* sequences displayed an identity ≥ 95% with a *Vibrio* type strains (see material and methods section "Population structure analysis").

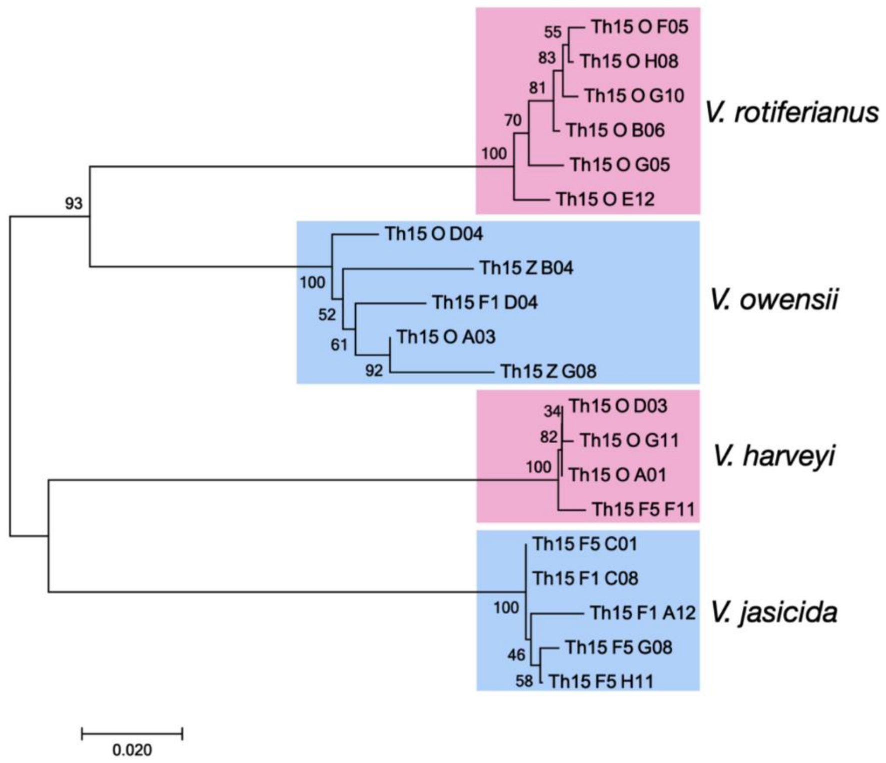

**Fig S3. Discrimination of Harveyi species by *rctB* sequencing.**

Phylogenetic tree constructed using the Neighbor-Joining method, the optimal tree with the sum of branch length = 0.49310882 is shown. The percentage of replicate trees in which the associated taxa clustered together in the bootstrap test (1000 replicates) are shown next to the branches. The tree is drawn to scale, with branch lengths in the same units as those of the evolutionary distances used to infer the phylogenetic tree. The evolutionary distances were computed using the Maximum Composite Likelihood method and are in the units of the number of base substitutions per site. All positions containing gaps and missing data were eliminated. There were a total of 494 positions in the final dataset. Evolutionary analyses were conducted in MEGA7.

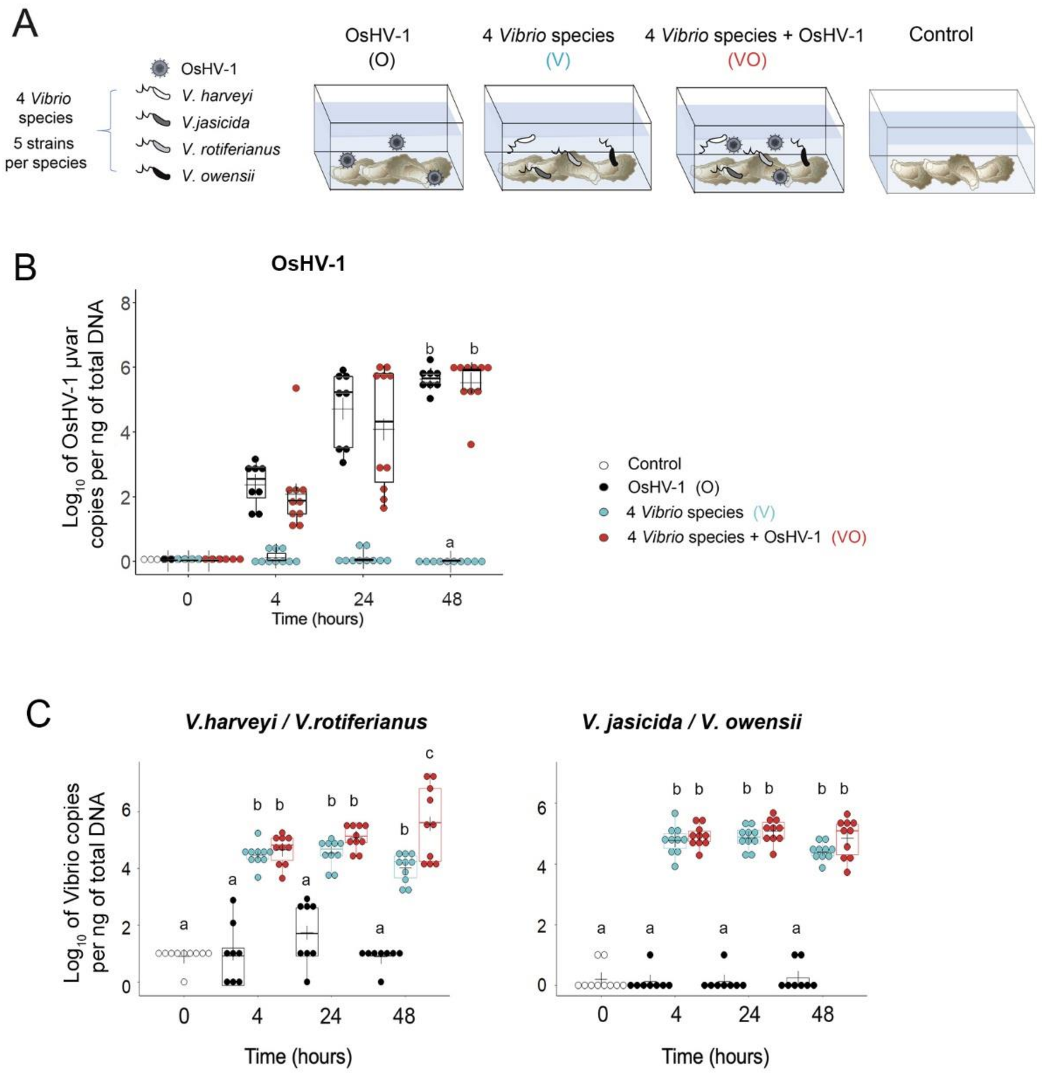

**Fig S4. OsHV-1 load in oysters exposed to the mesocosm experiment (Fig. 2)**

**(A) Design of a mesocosm experiment to recapitulate the complexity of the disease.** Oysters were immersed at 20°C in seawater containing OsHV-1 (O), 4 species of the Harveyi clade (V; *V. harveyi, V. rotiferianus, V. owensii, V. jasicida*), or both (VO). Oyster mortalities were monitored daily over 6 days. 10 individuals were sampled at each time point (0, 4, 24 and 48 h) for monitoring pathogens. **(B) Total OsHV-1 µVar quantified on individual oysters by qPCR.** Number of copies of OsHV-1 per ng of total DNA were $\log_{10}(x+1)$ transformed. Final time points were compared by a Kruskal-Wallis test. Significant differences between conditions are indicated with different letters ($p < 0.001$). **(C) qPCR monitoring of species of the Harveyi clade** on individual oysters using primer pairs discriminating *V. harveyi/V. rotiferianus* from *V. jasicida/V. owensii* (hrF1/R1 and oj-F1/R1, targeting a specific chemotaxis protein and *ompA* gene, respectively). Data show that OsHV-1 significantly enhanced colonization by *V. harveyi/V. rotiferianus* (VO) at 48h (mutiple t test, $p < 0.01$). No significant effect of OsHV-1 was observed on *V. jasicida/V. owensii* colonization. Each dot represents an individual.

A

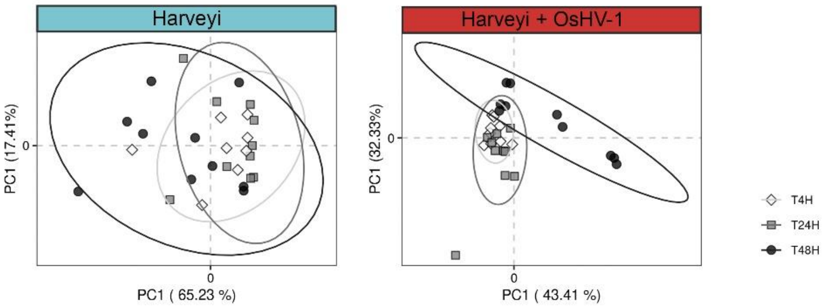

B

| Source of variation | DF | Sum Sq | Mean Sq | F | Pr (>F) |
|---|---|---|---|---|---|
| Harveyi.Time-steps | 2 | 0.22743 | 0.15190 | 2.2389 | 0.059 |
| Residual | 25 | 1.26975 | 0.84810 | | |
| Total | 27 | 1.49718 | 1.00000 | | |
| **Harveyi+OsHV-1.Time-steps** | 2 | 0.75804 | 0.29006 | 5.3113 | 0.001*** |
| Residual | 26 | 1.85537 | 0.70994 | | |
| Total | 28 | 2.61341 | 1.00000 | | |

**Fig S5. Monitoring of the dysbiosis by *rctB*-metabarcoding**

The colonization of oysters by the *Vibrio* Harveyi community in a mesocosm experiment (see Fig. 2) was monitored trough an *rctB* gene-based barcoding. **(A) Principal Component Analysis (PCoA) ordination plots of Bray-Curtis dissimilarities for the *Vibrio* Harveyi community.** PCoA results are presented for each experimental condition. Left panel corresponds to animals exposed to 4 species of the Harveyi clade (V; *V. harveyi, V. rotiferianus, V. owensii, V. jasicida*. Right panels corresponds to oysters exposed simultaneously to the same 4 species of the Harveyi clade and OsHV-1. Each dot corresponds to the Harveyi community of one oyster. Different symbols are used for different time-steps (i.e. 4, 24 and 48 h). Ellipses represent the 95% confidence intervals for each group. Permutational multivariate analysis of variance (PERMANOVA) between the two experimental conditions and the three time-steps indicates significant differences Pr (>F) = 0.001. Data highlight the destabilization of the Harveyi community at 48h in the presence of both OsHV-1 µVar and *Vibrio* (right panel). No significant shift was observed at any time point when oysters were exposed to *Vibrio* only (left panel). **(B) Statistical analyses supporting Harveyi community destabilization at 48h in presence of OsHV-1.** Results of permutational multivariate analysis of variance (PERMANOVA) with 1,000 permutations using the *adonis2* function implemented in the vegan R package.

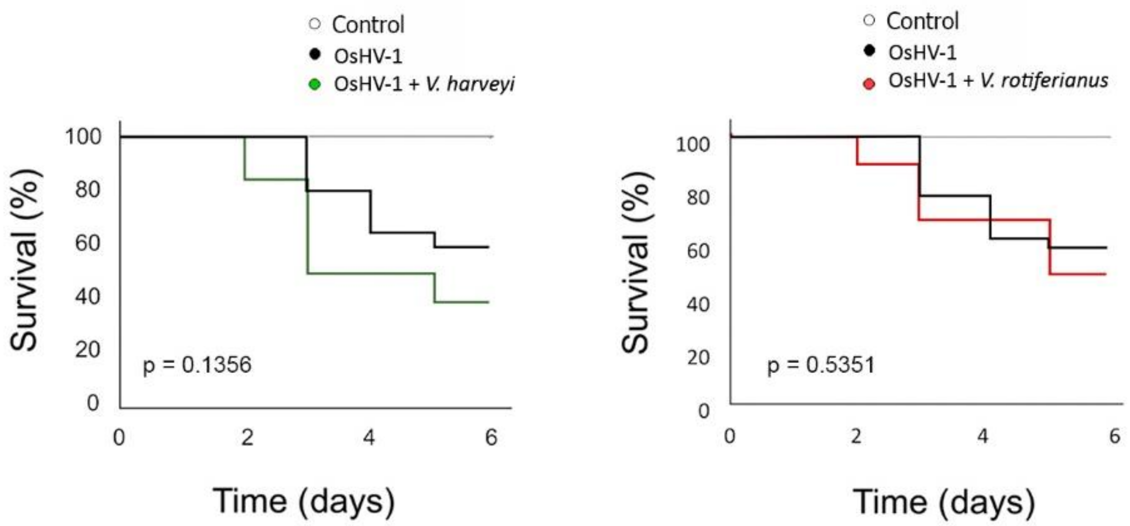

**Fig S6. Mortalities of oysters co-infected with OsHV-1 and single Vibrio species**

Kaplan-Meier survival curves were generated for oysters exposed to both *V. harveyi* Th15_O_G11 and OsHV-1 (green) or both *V. rotiferianus* Th15_O_G05 and OsHV-1 (red) compared to OsHV-1 only (black). No significant difference was observed between conditions (log-rank test, $p = 0.1356$ and $p = 0.5351$ for co-infections with *V. harveyi* and *V. rotiferianus*, respectively).

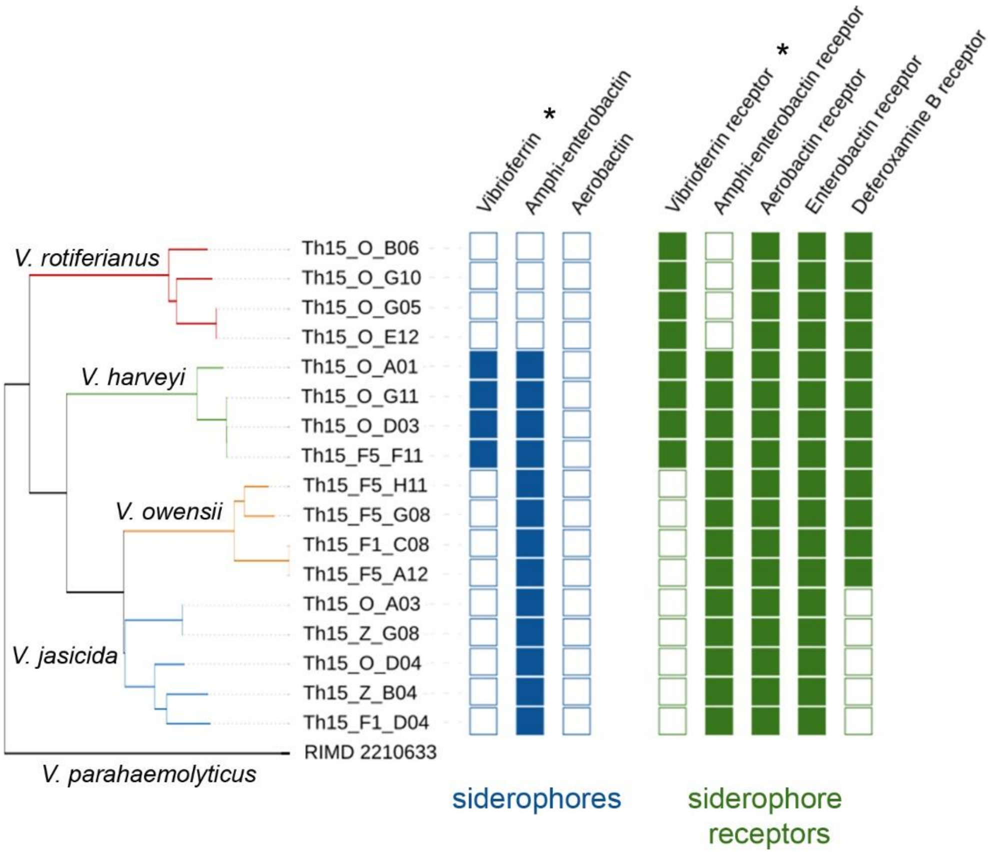

**Fig S7. Siderophores and siderophore uptake systems from Harveyi-related strains**

Comparative genomics between strains of *V. harveyi* (Th15_O_G11, Th15_F5_F11, Th15_O_D03, Th15_O_A01) and *V. rotiferianus* (Th15_O_G05, Th15_O_G10, Th15_O_B06, Th15_O_E12) point to a possible cheating behavior for siderophores in *V. rotiferianus,* which has receptors for at least 4 siderophores but does not seem to produce any. An asterisk highlights the Vibrioferrin biosynthetic pathway and uptake machinery.

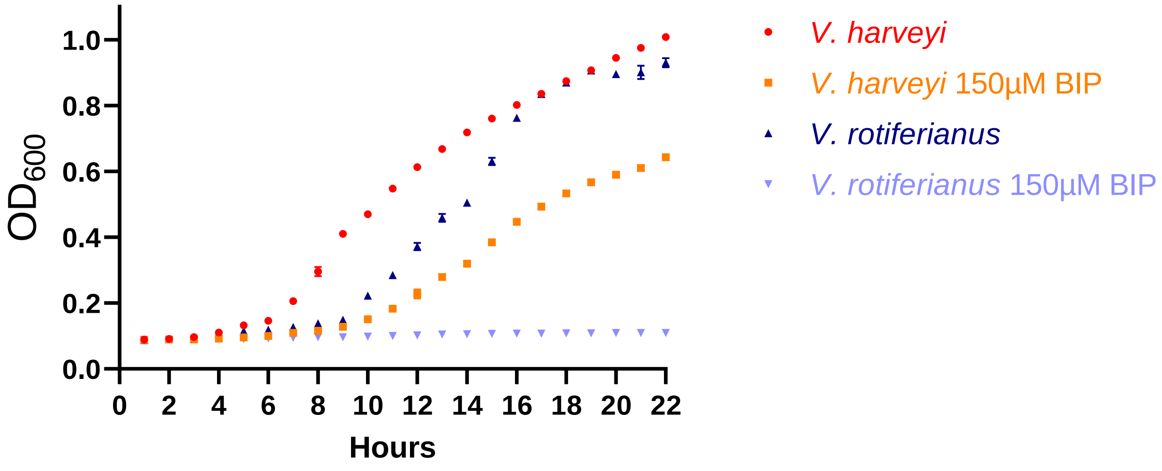

**Fig S8. Growth of *V. harveyi* and *V. rotiferianus* in iron-depleted conditions.**

Strains *V. harveyi* Th15_O_G11 and *V. rotiferianus* Th15_O_G05 were grown in Lewis water 1X supplemented with casaminoacids and Vitamin B. Iron-depletion was obtained by adding 150 µM 2,2'-bipyridine (BIP) to the minimal culture medium. Only *V. harveyi* could grow in such culture conditions.

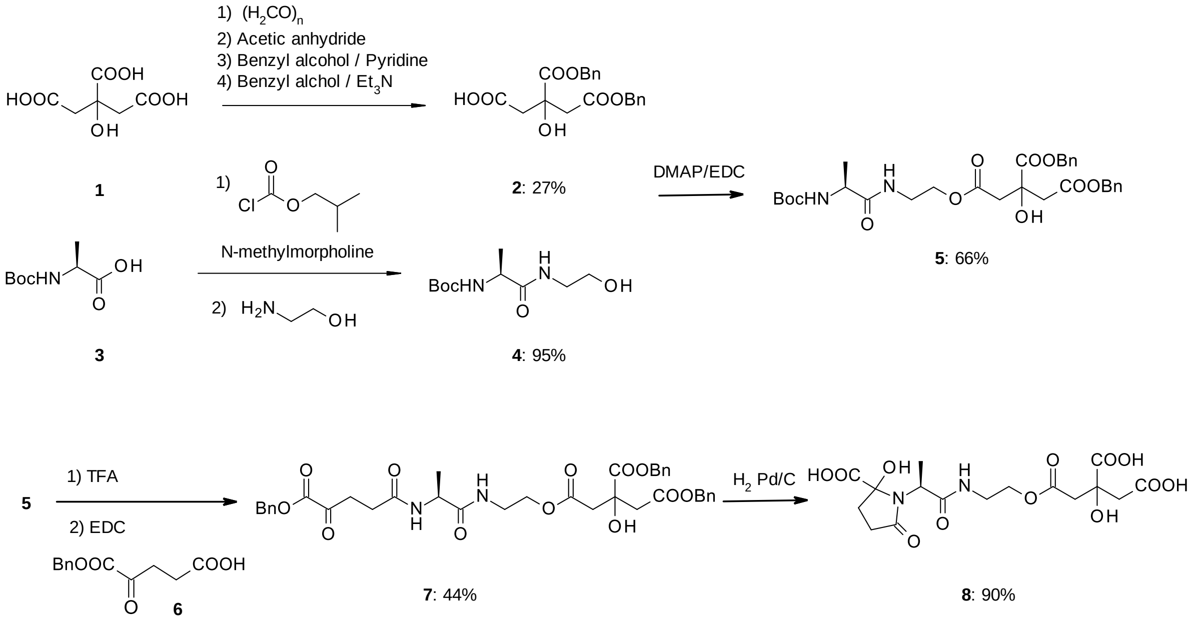

**Fig S9. Synthetic Vibrioferrin production following Takeuchi procedure [11].**

First, racemic dibenzylcitrate 2 was prepared starting from citrate 1 in four steps with an overall yield of 27% [12]. For the purpose of the present study optical resolution of 2 was considered unnecessary as we anticipated that the biological activity of racemic vibrioferrin 8 should be conserved. Subsequent Steglich esterification of dibenzylcitrate 2 by compound 4, which results from a reaction between N-Boc-alanine 3 and 2-aminoethanol, delivered compound 5. Acid mediated deprotection of compound 5 and coupling with benzyl ketogutaric acid 6 leads to compound 7 that was submitted to hydrogenolysis to afford vibrioferrin 8. Following this procedure, the overall yield from citrate 1 is 7% and 30 mg of vibrioferrin 8 were synthesized.

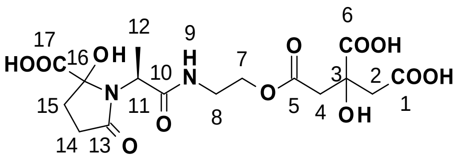

$^{1}H$ NMR ($D_2O$): [1.39 (d, J=6.9 Hz), 1.43 (d, J=7.5 Hz), 3H], 2.17 (m, 1H), 2.49-2.56 (m, 3H), [2.82 (d, J=16 Hz), 2.85 (d, J=12.0 Hz), 2H], 3.03 (d, J=16.0 Hz), 3.46 (m, 2H), 4.03 (m, 1H), 4.17 (m, 2H)

$^{13}C$ NMR ($D_2O$): 13.2,14.0 (C12), 29.1, 29.3 (C15), 32.0, 32.4 (C14), 38.3, 38.4 (C8), 43.3, 43.6 (C2-C4), 51.7, 51.8 (C11), 63.7, 63.8 (C7), 73.3 (C3), 91.1, 92.2 (C16), 171.2, 173.0, 173.4, 175.5, 176.6, 177.6 (C1, C5, C6, C10, C13, C17)

HRMS: Calculated for $C_{16}H_{22}N_2O_{12}$ $[M-H_2O]^+$: 417.1145, $[M+H]^+$: 435.1251, $[2M+H]^+$: 869.2424 found for $[M-H_2O]^+$: 417.1126 $[M+H]^+$: 435.1229 $[2M+H]^+$: 869.2392

$^{1}H$ NMR spectra of vibrioferrin **8**

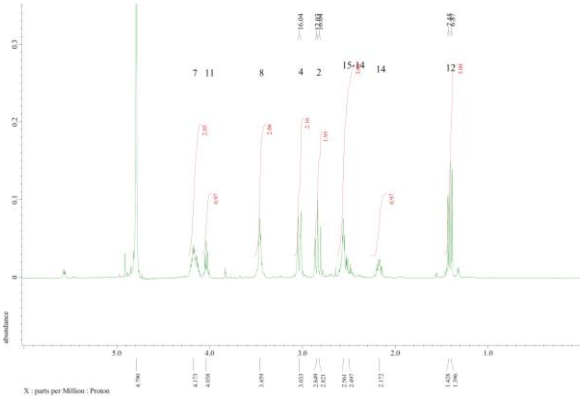

$^{13}C$ NMR spectra of vibrioferrin **8**

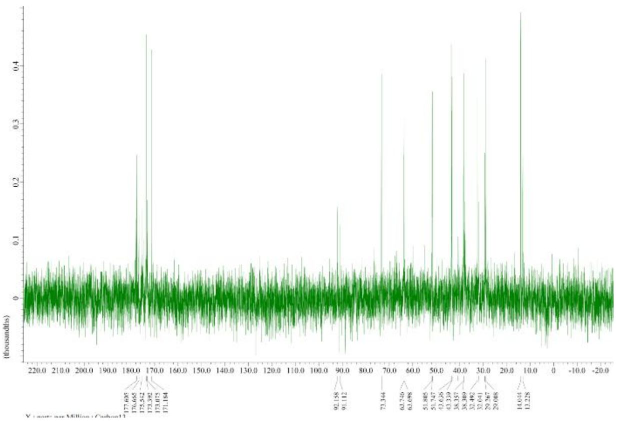

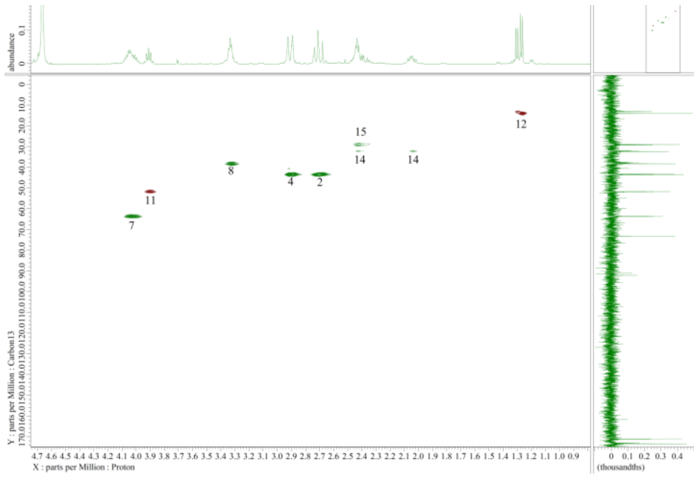

HRMS spectra of vibrioferrin **8**

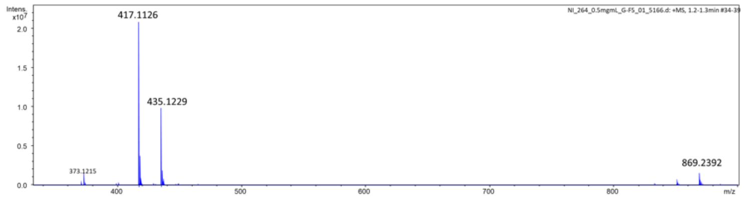

**Fig S10. Characterization of racemic vibrioferrin 8.**

NMR spectra were recorded on a JEOL ECZ-500R spectrometer, and the HRMS spectra on a Brucker Q-Tof maXis. The NMR or mass spectroscopy data are in good agreement with data from Takeushi et al [11]. The chemical shifts for the $^{1}H$ and $^{13}C$ NMR spectra are identified using the numbering provided with the vibrioferrin formula. The presented data are sufficient to ascertain that the synthesis of vibrioferrin was successfully completed.

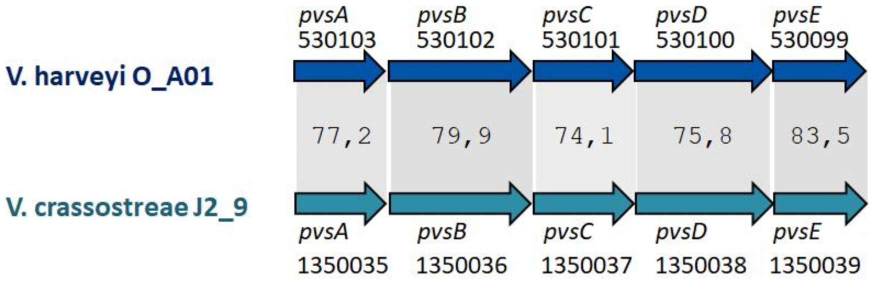

**Fig S11. Conservation of vibrioferrin biosynthesis pathway in *V. crassostreae***
Synteny alignment of predicted proteins implicated in vibrioferrin biosynthesis pathway in *V. harveyi* O_A01 and *V. crassostreae* J2_9 genomes. Gene annotations and CDS reference are indicated. Identity between homologous proteins are indicated (in %) and represented by grey shades. Protein identities were calculated using BLOSUM62 comparison matrix and SIM Alignment tool for protein sequences (https://web.expasy.org/sim/)

### Supplementary Materials and Methods

#### Mutagenesis and mutant validation

*E. coli* strains used for mutagenesis (Table S4) were routinely grown at 37°C in Luria-Bertani (LB) medium and *V. rotiferianus* Th15_O_G05 was cultivated at 20°C in LB supplemented with 20 g/l (final concentration) NaCl (LBS). If necessary antibiotics and components were added to the media: kanamycin (Km, 100 µg/ml), chloramphenicol (Cm, 5 µg/ml), D-glucose (D-Glc, 0.3 g/l), L-arabinose (L-ara, 0.2%), diaminopimelic acid (DAP, 0.3 mM) and agar (15 g/l).

The DNA fragment containing *pvuA1* and *pvuA2* was deleted in *V. rotiferianus* Th15_O_G05 by double homologous recombination between a suicide plasmid (Table S5) and the chromosome. Briefly, two fragments of around 800 bp located upstream (up) of the *pvuA1* gene (THOG05_v1_100041) and downstream (down) of the *pvuA2* gene (THOG05_v1_100042) were synthetized and assembled by GeneArt (Invitrogen) and supplied into a pMKRQ plasmid. The synthetic up-down fragment was released from pMKRQ using *Xho*I and *Eco*RI and ligated into the pLP12 plasmid (Luo *et al.*, 2015) (digested with same enzymes) using T4 DNA ligase (Promega) according to manufacturer recommendations. The ligation mixture was transformed by heat shock into competent cells of *E. coli* DH5α λpir. After PCR-confirmation, a clone containing the pLP12 in which the up-down fragment was inserted (named pAM010) was stored in 20% glycerol at -80°C. The purified pAM010 was transferred by heat shock into competent cells of *E. coli* β3914 [6] carrying a RP4 origin of transfer. Finally, pAM010 was transferred to *V. rotiferianus* Th15_O_G05 by a triparental mating procedure, adapting a recently published method [7]. Briefly, after overnight growth, 200 µl of the *E.coli* β3914 donor strain carrying the pAM010 were centrifuged, 200 µl of the *E. coli* GEB883 helper strain carrying the pEVS104 plasmid [13] were added to the pellet, centrifuged, and 800 µl of the *V. rotiferianus* Th15_O_G05 were added to the pellet and centrifuged. The cell pellet was resuspended with 20 µl of LBS+DAP and spotted on a 0.2 µm acetate filter (Sartorius Stedim biotech) placed on the LBS+DAP plate. This plate was then incubated at 20°C for 24 h. The filter was rinsed with 1 ml filter-sterilized seawater (FSSW) and the suspension cell was centrifuged. The pellet was resuspended with 50 µl of LBS+DAP, plated on LBS+D-Glc+Cm plate and then incubated at 20°C for 24 h to 48 h. The obtained colonies were restreaked on LBS+D-Glc+Cm and the integration of pAM010 in the chromosome of *V. rotiferianus* Th15_O_G05 was determined by PCR, using Insert-pLP12-R and del-pvuA1-A2-OG05-F primers. Several colonies with an integration of the suicide plasmid in the "upstream" or in the "downstream" region of the to-be-deleted DNA fragment were grown overnight in LBS at 20°C. Cultures were serially diluted and 100 µl were plated on LBS+L-ara to the excision of the pAM010. After a 24 h-incubation at 20°C, colonies were restreaked on LBS+Cm and LBS+L-ara to control the Cm-sensitivity. The Cm-sensitive clones were tested for deletion of *pvuA1* and *pvuA2* genes by using primers del-pvuA1-A2-OG05-F and del-pvuA1-A2-OG05-R (Table S5). A *V. rotiferianus* Th15_O_G05 mutant strain deleted for the *pvuA1-2* genes was stored in glycerol at -80°C (strain *V. rotiferianus* Th15_O_G05 Δ*pvuA1-2*).

#### Standard curves for absolute qPCR quantification

Standard curves of known concentration of *Vibrio* genomes were generated using the relation between the concentration of DNA and the theoretical copy number of genomes, calculated on the basis of the DNA mass divided by molecular weight of the genome. They were also validated by the limit dilution method, assuming that the dilution at which 1 replicate in 10 was positive corresponds to 1 copy.
